# The temporal refinement of *Dach1* is a key step in the functional maturation of primary somatosensory neurons

**DOI:** 10.1101/2024.07.18.604081

**Authors:** Tünde Szemes, Alba Sabaté San José, Abdulkader Azouz, Maren Sitte, Gabriela Salinas, Younes Achouri, Sadia Kricha, Laurence Ris, Kristy Red-Horse, Eric J. Bellefroid, Simon Desiderio

## Abstract

During somatosensory neurogenesis, neurons are born in an unspecialized transcriptional state, with several transcription factors following a broad-to-restricted expression dynamic as development proceeds, supporting neuron subtype identities. The relevance of this temporal refinement remains however unclear, these broad-to-restricted transcription factors being selectively involved in neurons in which they are ultimately maintained. Here we found that *Dach1* encodes for a *bona fide* broad-to-restricted transcription factor retained and required in tactile somatosensory neurons. Within developing nociceptors, we demonstrate that Prdm12 contributes to *Dach1* extinction. Using genetic approaches to prevent its temporal restriction during somatosensory development, we reveal that *Dach1* refinement is a prerequisite for the appropriate transcriptional maturation of somatosensory subtypes from which it becomes ultimately excluded. These findings highlight the essential role played by Dach1 during somatosensory neuron development. They further demonstrate that the broad-to-restricted temporal pattern followed by several transcription factors is physiologically relevant to achieve appropriate transcriptional maturation of somatosensory neurons.

## Introduction

Primary somatosensory neurons constitute a heterogenous population of peripheral nervous system (PNS) neurons tuned to respond to dedicated stimuli emanating from the external and internal environment. These neurons are found in somatosensory ganglia such as the trigeminal ganglia innervating the head and the dorsal root ganglia (DRG) innervating the body. They support the broad spectrum of somesthesia and are segregated based on their conduction velocity and degree of myelination (Aα-, Aβ-, Aδ- or C subtypes, from heavily myelinated to unmyelinated), their morphology and their transcriptional and physiological features. They can be broadly categorized into three cardinal functional populations; the proprioceptive neurons that innervate tendons and muscles to provide a sense of body position; the low-threshold mechanoreceptors (LTMR) which comprise neurons of C-, Aδ- or Aβ-subtypes involved in touch through the detection of innocuous mechanical stimuli, and the nociceptors comprising a set of largely functionally polymodal neurons of Aδ- or C-subtype involved in nociception, detection of temperature or pruriception^1^.

In the body, these neurons stem from a pool of neural crest cells biased to the somatosensory lineage through the transient expression of the proneural factors Neurog1 and Neurog2^1–3^. The segregated expression of dedicated neurotrophin receptors later allows a target-derived signal-based crosstalk with the environment to support the survival and maturation of these neurons and drive some aspects of their subtype-specific morphological and transcriptional features^3–5^. Thereby, developing nociceptors initially express the neurotrophin receptor TrkA and later diversify into peptidergic or non-peptidergic nociceptors which respectively maintain TrkA or express Ret^6^ while proprioceptors express TrkC and A-LTMR subtypes rely on selective or combinatorial expression of Ret, TrkB or TrkC^3,5^. A general view of neuronal development is that neurons mature through the orderly temporal induction and action of stepwise transcriptional programs^7^. Hence, several subtype-restricted transcription factors (TFs) are also widely involved in the development and maturation of these somatosensory neurons at defined periods during their maturation process^2,3,8^.

While case by case functional studies have proven valuable to understand their selective developmental requirement in somatosensory subtypes, recent findings highlighted that a cohort of these subtype-restricted TF are initially widely co-expressed in newborn somatosensory neurons, following extinction of *Neurog1* and *Neurog2*^9^. Newborn somatosensory neurons thus transition from an unspecialized state characterized by wide co-expression of TFs to subsequent stages characterized by the resolution of restricted combinatorial expression of these TFs that transcriptionally define and specify somatosensory subtype identities^1,9,10^. Loss of function studies aimed at identifying the role of genes such as *Pou4f2, Pou4f3, Runx1, Runx3* or *Shox2* which encode for some of these broad-to-restricted TFs have demonstrated that their depletion from the time of their broad expression period results in defects selectively compartmentalized to the somatosensory subtypes in which they are ultimately maintained^9,11–15^. These observations raise the question of the biological significance of this broad-to-restricted temporal pattern as TFs showing such a developmental dynamic do not appear to play prominent function at the time of their broad expression period.

Searching to describe new TFs with broad-to-restricted expression in developing DRG, we identified the gene encoding for the helix-turn-helix transcription factor Dach1, which relates to the sno/ski family of corepressors^16^. We found that Dach1 is initially ubiquitously expressed in unspecialized postmitotic somatosensory neurons and becomes gradually selectively enriched in TrkB-expressing A-LTMR neurons as development proceeds. In the nociceptive lineage, we demonstrate that this concomitant temporal extinction of Dach1 involves the nociceptor-specific transcriptional regulator Prdm12. Genetic depletion of *Dach1* during its broad or restricted expression period results in A-LTMR restricted defects, thus classifying Dach1 as a *bona fide* broad-to-restricted TF. Gain-of-function strategies to prevent the temporal refinement of Dach1 result in wide transcriptional defects in neurons that should repress Dach1, functionally reflected at adulthood by reduced nociception. Altogether, these results establish Dach1 as a new component of the transcriptional pathways leading to the maturation of somatosensory neurons, for which both the silencing and maintenance of its expression are critical. They further provide clues to the biological significance of the tight temporal regulation of broad-to-restricted TFs as a key event to achieve proper development of somatosensory neurons.

## Results

### Dach1 is expressed during somatosensory neurons development following a broad-to-restricted expression pattern

Recent advances in the understanding of the transcriptional maturation and diversification of developing and adult somatosensory neurons arose from newly generated single-cell transcriptomic atlases. These databases can be leveraged to highlight uncharacterized genes selectively expressed in discrete somatosensory subtypes^9,17,18^. Looking for genes encoding transcriptional regulators with such discrete pattern, we identified *Dach1*, a member of the sno/ski family of corepressors, which is putatively enriched at adulthood in myelinated LTMR neurons **(Figure S1A)**.

Immunostainings were performed on murine embryonic DRG sections to define Dach1 expression during somatosensory neuron development. From E10.5 onward, Dach1 is detected from Sox10^+^ somatosensory progenitors and becomes strongly established as these cells become postmitotic and lose Sox10 **(Figure 1A)**. Between E10.5 and E18.5, comparison of Dach1 and Islet1, a general marker of postmitotic somatosensory neurons, indicated that while most neurons retain Dach1 at early developmental stages, it becomes widely downregulated at E14.5 (**Figure 1B-C**). Analysis of the fluorescence level of Dach1 immunostaining as a readout of its expression in Islet1^+^ neurons revealed that Dach1 refinement occurs around E13.5, corresponding to the end of the period of overt neurogenesis in DRG^1,2^ **(Figure 1D)**. To determine the identity of neurons maintaining a strong Dach1 staining at E18.5, double immunostainings were carried out with Prdm12 labelling nociceptors and C-LTMR, TrkB labelling A-LTMR, TrkC labelling A-LTMR and proprioceptors and Parvalbumin (Parvalb) labelling proprioceptors. 85,09% ± 1,69% (Mean ± S.D., n = 2) and 16,34% ± 2,40% of E18.5 Dach1^+^ neurons respectively colocalized with TrkB and TrkC. Only a negligible proportion of Dach1*^+^* neurons colocalized with Prdm12 (3,90% ± 0,18%) or Parvalbumin (1,97% ± 0,31%). These results demonstrated that Dach1 becomes enriched in TrkB-expressing A-LTMR. This was further confirmed by monitoring the co-expression of Dach1 with Ret, MafA and c-Maf, other known markers of A-LTMR subpopulations **(Figure 1E)**.

**Figure 1.**
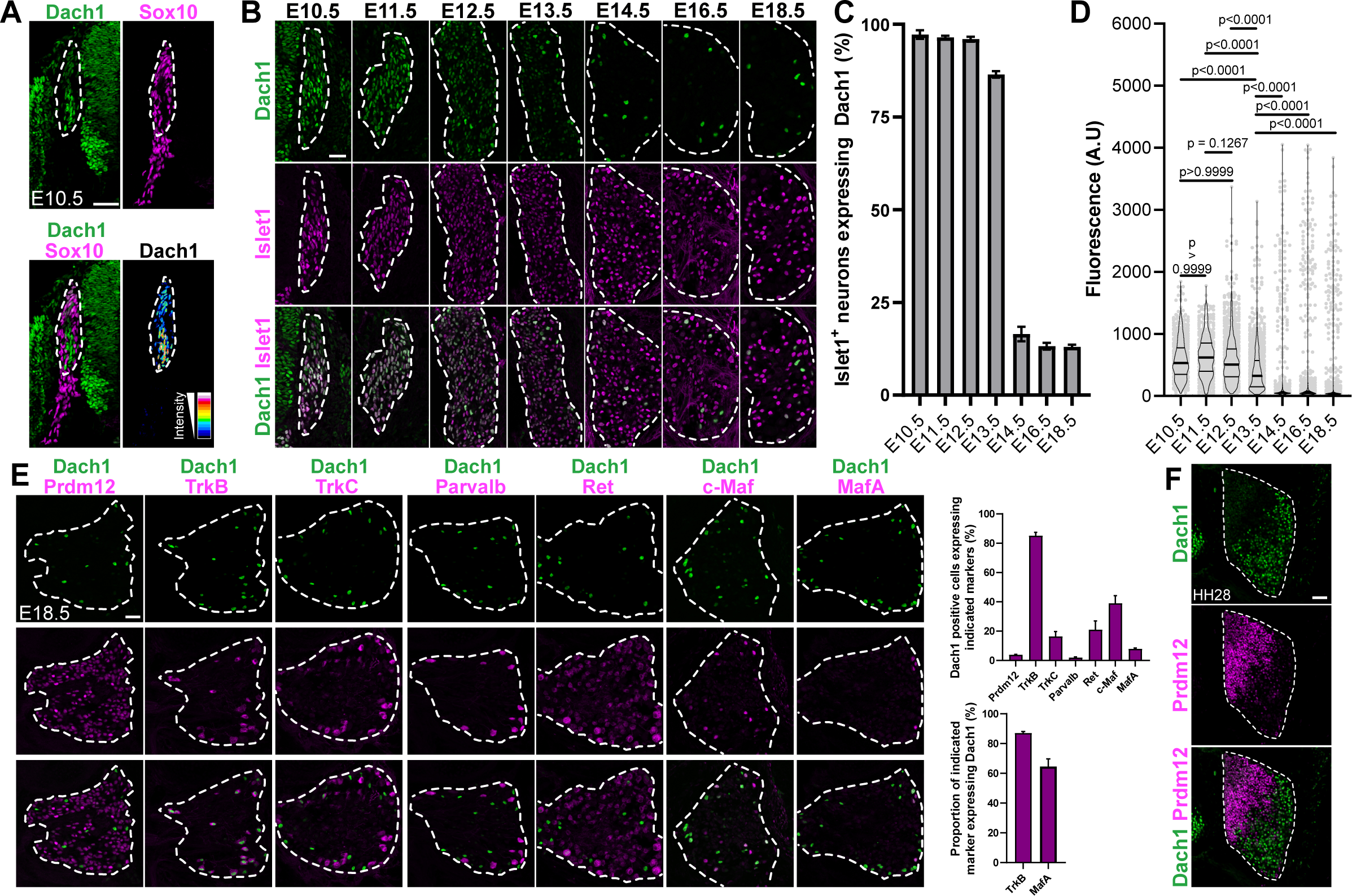
Dach1 follows a broad-to-restricted expression pattern in developing dorsal root ganglia. **(A)** Double immunostaining targeting Dach1 and Sox10 performed on transverse section of E10.5 murine dorsal root ganglion (DRG) at the thoracic level. The lower right panel shows a fluorescence intensity pseudo-coloured image of Dach1 staining to highlight its increased expression in Sox10-negative DRG neurons. **(B)** Representative pictures of double immunostainings of Dach1 with the pan-neuronal marker Islet1 performed on thoracic DRG sections of murine embryos at the indicated developmental stages. **(C)** Histogram representing the mean percentage of Islet1^+^ neurons co-expressing Dach1 in DRG transverse sections at indicated developmental stages. Data represent the mean ± SEM of quantifications from two embryos per stage. **(D)** Quantification of Dach1 fluorescence staining (arbitrary unit, A.U.) among Islet1^+^ DRG neurons at indicated developmental stages. Each grey point represents the fluorescence level of Dach1 in a single Islet1^+^ nucleus. The total number of Islet1^+^ neurons measured at each stage were: 611 at E10.5, 489 at E11.5, 1145 at E12.5, 1186 at E13.5, 914 at E14.5, 998 at E16.5 and 821 at E18.5. Overlayed violin plots represent data distribution with first and third quartiles indicated by plain lines and median indicated by thick plain line. Kruskal-Wallis test with Dunn’s post hoc multiple comparison. **(E)** Left, double immunostainings with Dach1 and indicated markers performed on E18.5 thoracic DRG transverse sections. Right, quantifications of the colocalization of Dach1 and indicated somatosensory subtype markers. Data are represented as mean ± SEM, at least two embryos analysed per condition. **(F)** Double immunostaining of Dach1 and Prdm12 on transverse DRG section of embryonic chicken at Hamburger & Hamilton (HH) stage 28. DRG in pictures are delineated by white dashed lines. Scale bars, 50 µm.

Contrary to what is observed in mice, chicken embryonic DRG show a clear spatial segregation of neuronal subtypes, with nociceptors and mechano/proprioceptors being respectively confined to the dorsomedial and ventrolateral part of the DRG^19^. Co-immunostaining of Prdm12 and Dach1 performed on HH28 chicken DRG sections revealed that Prdm12^+^ and Dach1^+^ cells are dorsomedially and ventrolaterally segregated as expected **(Figure 1F)**. Analysis of Dach1 expression in adult murine DRG showed the maintenance of its preferential localization to TrkB-expressing neurons compared to other cardinal subtypes. Single-cell RNA-seq databases indicate that *Dach1* is notably enriched in TrkB^+^ Aδ-LTMR. This observation was further confirmed *in situ* by analysing the colocalization of Dach1 with the Aδ-LTMR marker *Colq*^9^ (**Figure S1B-C)**. These findings demonstrate that Dach1 follows a broad-to-restricted expression pattern in developing DRG to become progressively compartmentalized to Aβ- and Aδ-LTMR.

### The nociceptor determinant Prdm12 contributes to the temporal refinement of Dach1

The temporal restriction of Dach1 to A-LTMR neurons implies its progressive repression by other somatosensory subtypes, including by nociceptors, which represent the vast majority of developing somatosensory neurons^2^. This temporal extinction was evidenced through co-immunostaining of Dach1 with Prdm12 and TrkA, two general markers of developing nociceptors, before (E12.5) and after (E16.5) its temporal refinement (**Figure 2A**). Previous reports suggest that neurotrophin-derived signalling pathways are involved in some aspects of the transcriptional refinement controlling subtype identity. More specifically, the NGF-TrkA signalling pathway plays a predominant role in the embryonic survival and development of the nociceptive lineage^6,9^.

**Figure 2.**
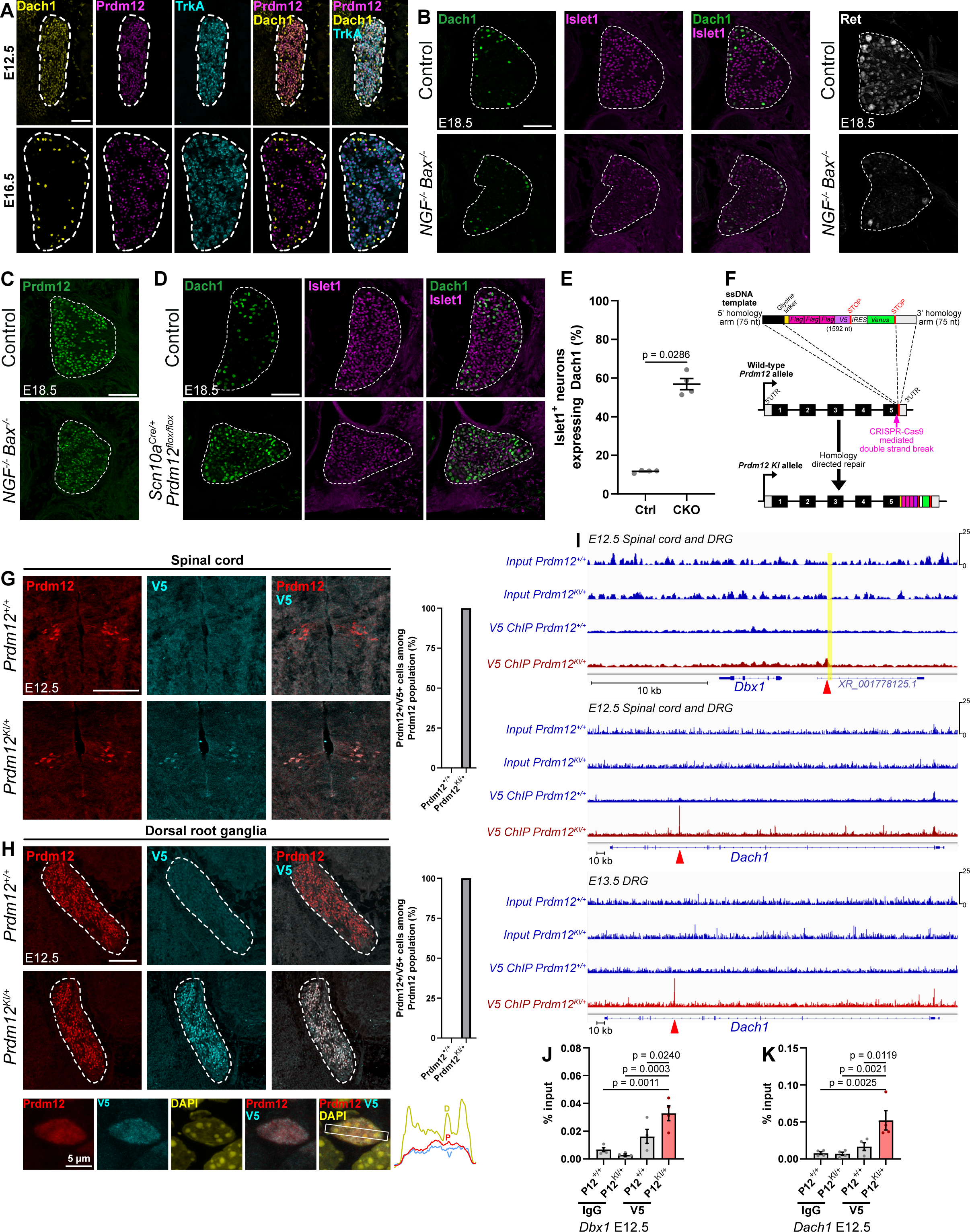
Prdm12 contributes to the temporal refinement of Dach1 independently of NGF signalling. **(A)** Triple immunostaining of Dach1, Prdm12 and TrkA performed on E12.5 and E16.5 DRG transverse sections of control embryos. **(B)** Left, double immunostaining of Dach1 and Islet1 performed on E18.5 DRG transverse sections of control and *NGF*^-/-^;*Bax*^-/-^ embryos. Note the maintenance of Dach1 restricted expression pattern in absence of *NGF*. Right, Immunostaining targeting Ret performed as a positive control showing the proper inactivation of the NGF signalling pathway in *NGF*^-/-^;*Bax*^-/-^ E18.5 DRG. **(C)** Immunostaining targeting Prdm12 performed on E18.5 DRG transverse sections of control and *NGF*^-/-^;*Bax*^-/-^ embryos. Note the maintained expression of Prdm12 in absence of NGF. **(D)** Double immunostaining of Dach1 and Islet1 performed on E18.5 DRG transverse sections of control and *Scn10a^Cre/+^; Prdm12^flox/flox^* embryos. Note the broadening of Dach1^+^ cells in absence of Prdm12. **(E)** Quantification analysis of the mean percentage of Islet1^+^ neurons expressing Dach1 in DRG of E18.5 control and *Scn10a^Cre/+^; Prdm12^flox/flox^* (CKO) embryos. Mann-Whitney tests. **(F)** Schematic representation of the CRISPR-Cas9-based strategy used to generate the *Prdm12^KI^* allele. **(G)** Left, double immunostaining of Prdm12 and V5 performed on E12.5 spinal cord transverse sections of *Prdm12^+/+^*and *Prdm12^KI/+^* embryos. Right, quantification analysis of the mean percentage of V5^+^ nuclei expressing Prdm12 in E12.5 spinal cords of *Prdm12^+/+^* and *Prdm12^KI/+^*embryos.**(H)** Left, double immunostaining of Prdm12 or TrkB with V5 performed on E12.5 DRG transverse sections of *Prdm12^+/+^* and *Prdm12^KI/+^* embryos or *Prdm12^KI/+^* embryos alone. Top right, quantification analysis of the mean percentage of V5^+^ nuclei expressing Prdm12 in E12.5 DRG of *Prdm12^+/+^* and *Prdm12^KI/+^* embryos. Bottom right, quantification analysis of the mean percentage of V5^+^ nuclei found in TrkB^+^ neurons in E12.5 DRG of *Prdm12^KI/+^* embryos. Bottom, colocalization analysis of Prdm12 and V5 in a E12.5 DRG cell counterstained with DAPI (scale bar, 5µm). **(I)** Mouse genomic region surrounding *Dbx1* (top) or *Dach1* (middle) showing sequencing data for input DNA recovered from *Prdm12^+/+^* and *Prdm12^KI/+^* and V5 ChIP-enriched DNA recovered from *Prdm12^+/+^* and *Prdm12^KI/+^* E12.5 spinal cord with attached DRG. Bottom trace view shows that the peak observed at E12.5 is still found following V5 targeted ChIP-seq performed on isolated E13.5 DRG. Peaks of interest are highlighted by red arrowheads. The homologous Prdm12 bound region near *Dbx1* previously characterized in *Xenopus laevis* is highlighted by a yellow band. **(J)** ChIP-qPCR validation of the peak identified in the vicinity of the *Dbx1* locus performed on dissected spinal cord and DRG of E12.5 embryos of indicated genotypes. Graphical data in this panel and in the subsequent ones are presented as mean ± SEM. One way ANOVA test with Dunnett’s post hoc multiple comparison. **(K)** ChIP-qPCR validation of the peak identified in the fourth intron of the *Dach1* gene performed on dissected spinal cord and DRG of E12.5 embryos of indicated genotypes. One way ANOVA test with Dunnett’s post hoc multiple comparison. DRG in pictures are delineated by white dashed lines. Scale bars, 100 µm.

As most neurons downregulating Dach1 are inferred to be developing nociceptors, we thus analysed Dach1 expression in DRG of *NGF^-/-^; Bax^-/-^* E18.5 embryos compared to control ones. While Ret expression was abolished in nociceptors of *NGF^-/-^; Bax^-/-^* mutants as previously reported^6^, Dach1 expression was not broadened, indicating that Dach1 temporal restriction does not depend on the NGF-TrkA signalling (**Figure 2B**). In DRG of *NGF^-/-^; Bax^-/-^* E18.5 embryos, we also analyzed the expression of the transcriptional regulator Prdm12, a critical determinant of the development of the nociceptors whose invalidation results in agenesis of the entire nociceptive lineage^20–23^. **Figure 2C** shows that Prdm12 is still expressed in DRG of *NGF^-/-^; Bax^-/-^* E18.5 embryos suggesting that its expression, like that of Dach1, is independent of NGF. Prdm12 is mainly known to act as a putative transcriptional repressor^24^. In developing DRG, its loss results in the ectopic expression of visceral neuron determinants in a portion of putative nociceptive precursors^20–22,25^. This prompted us to hypothesize that it may be involved in Dach1 refinement. However, given the combination of rapid cell loss and ectopic Phox2a/b expression in newborn nociceptors constitutively lacking *Prdm12*, unambiguous analysis of Dach1 refinement in DRG of *Prdm12* KO embryos is challenging. To overcome this difficulty, we generated *Scn10a^Cre/+^; Prdm12^flox/flox^* (*Prdm12* cKO) mice which result in late post-mitotic embryonic depletion of Prdm12 in developing nociceptors without cell loss or ectopic expression of Phox2a/b^25^. Analysis of DRG of E18.5 *Prdm12* cKO embryos revealed broadened expression of Dach1 (**Figure 2D-E**).

To further investigate the mechanism involving *Dach1* repression by Prdm12, we sought to perform chromatin immunoprecipitation (ChIP) experiments to determine whether *Dach1* may represent a direct transcriptional target of Prdm12. As no ChIP-grade Prdm12 antibodies have been described so far, we generated a Prdm12 knock-in (KI) mouse line in which Flag and V5 coding sequences were inserted in frame downstream of the endogenous *Prdm12* coding sequence, resulting in the production of a Prdm12 protein fused to three Flag and one V5 tags in its C-terminal region (*Prdm12^KI^*, **Figure 2F**). *Prdm12^KI/KI^* mice were viable, fertile and raised to adulthood without obvious abnormalities contrary to the birth lethal phenotype of *Prdm12^-/-^*mice^20^, indicating that addition of the tags does not preclude Prdm12 function. In the developing nervous system, *Prdm12* is notably found in spinal cord p1 progenitors^24^. Accordingly, immunostainings targeting Prdm12 and V5 epitopes performed on E12.5 *Prdm12^KI/+^* spinal cord and DRG transverse sections indicated that V5 faithfully recapitulates Prdm12 expression in both structures **(Figure 2G-H)**.

ChIP was subsequently applied to *Prdm12^KI/+^* and *Prdm12^+/+^* E12.5 isolated spinal cords with attached DRG or E13.5 isolated DRG using a V5 antibody and next generation sequencing was performed on harvested DNA to identify Prdm12 bound regions. A previous study reported that in *Xenopus laevis* animal cap explants treated with retinoic acid and overexpressing *Noggin* and *Prdm12*, Prdm12 binds downstream of the homeobox gene *Dbx1*, one of its repressed targets during spinal cord neurogenesis^24^. Mapping this genomic bound region in the resulting dataset to the murine genome, we consistently observed a specific enrichment of Prdm12 binding in a genomic region downstream of the murine *Dbx1* locus that is homologous to the one previously reported in *Xenopus laevis* (**Figure 2I**). This result was further validated by ChIP-qPCR, confirming that our ChIP-seq successfully identified Prdm12 bound genomic regions (**Figure 2J**). Analysis of the ChIP-seq dataset around the *Dach1* locus highlighted an intronic Prdm12 binding site in the *Dach1* gene both in the E12.5 spinal cord with associated DRG dataset and in the E13.5 isolated DRG dataset, indicating that the binding indeed occurs in somatosensory neurons (**Figure 2I**). Binding to this putative regulatory region was further validated by ChIP-qPCR (**Figure 2K**). Together, these results indicate that Prdm12 contributes to the temporal restriction of *Dach1* in the nociceptive lineage, independently of the NGF pathway.

### Loss of Dach1 results in tactile defects without affecting somatosensory neurogenesis

To evaluate the involvement of *Dach1* in somatosensory neuron development, we relied on two strategies aiming at depleting *Dach1* during (I) its early broad expression period or (II) its A-LTMR restricted expression period. To achieve this purpose, mice carrying a *Dach1* floxed exon 2 (*Dach1^flox/flox^*) were crossed with *Wnt1^Cre^*or *Advillin^Cre^* mice to respectively deplete *Dach1* from the neural crest or from postmitotic somatosensory neurons^26–28^. While *Advillin^Cre/+^; Dach1^flox/flox^* conditional knockout (*AD1* cKO) mice are viable and fertile, *Wnt1^Cre/+^; Dach1^flox/flox^* (*WD1* cKO) mice die soon after birth, suggesting that *Dach1*-associated neural crest defects contribute to the postnatal lethality previously reported in *Dach1* constitutive knockouts^29^. Immunostaining of Dach1 confirmed its extinction in both lines by E18.5 (**Figure S2A**). Furthermore, Dach1 was already abolished by E12.5 in DRG of *WD1* cKO embryos, indicating that its depletion occurs as expected during its pan-sensory period (**Figure S2B**). In contrast, *Dach1* depletion in *AD1* cKO embryos became effective around E14.5, when Dach1 is predominantly restricted to TrkB^+^ A-LTMR (**Figure S2C-E**). In *WD1* cKO and *AD1* cKO E18.5 embryos, the counting of Islet1^+^ somatosensory neurons in thoracic DRG revealed no difference between mutants and controls. We also examined *bona fide* general markers of somatosensory subtypes and found again no difference between the cohorts. Overall, these results show that depletion of Dach1 does not impact neurogenesis and does not alter the proportions of the main neuron subtypes (**Figure 3A**).

**Figure 3.**
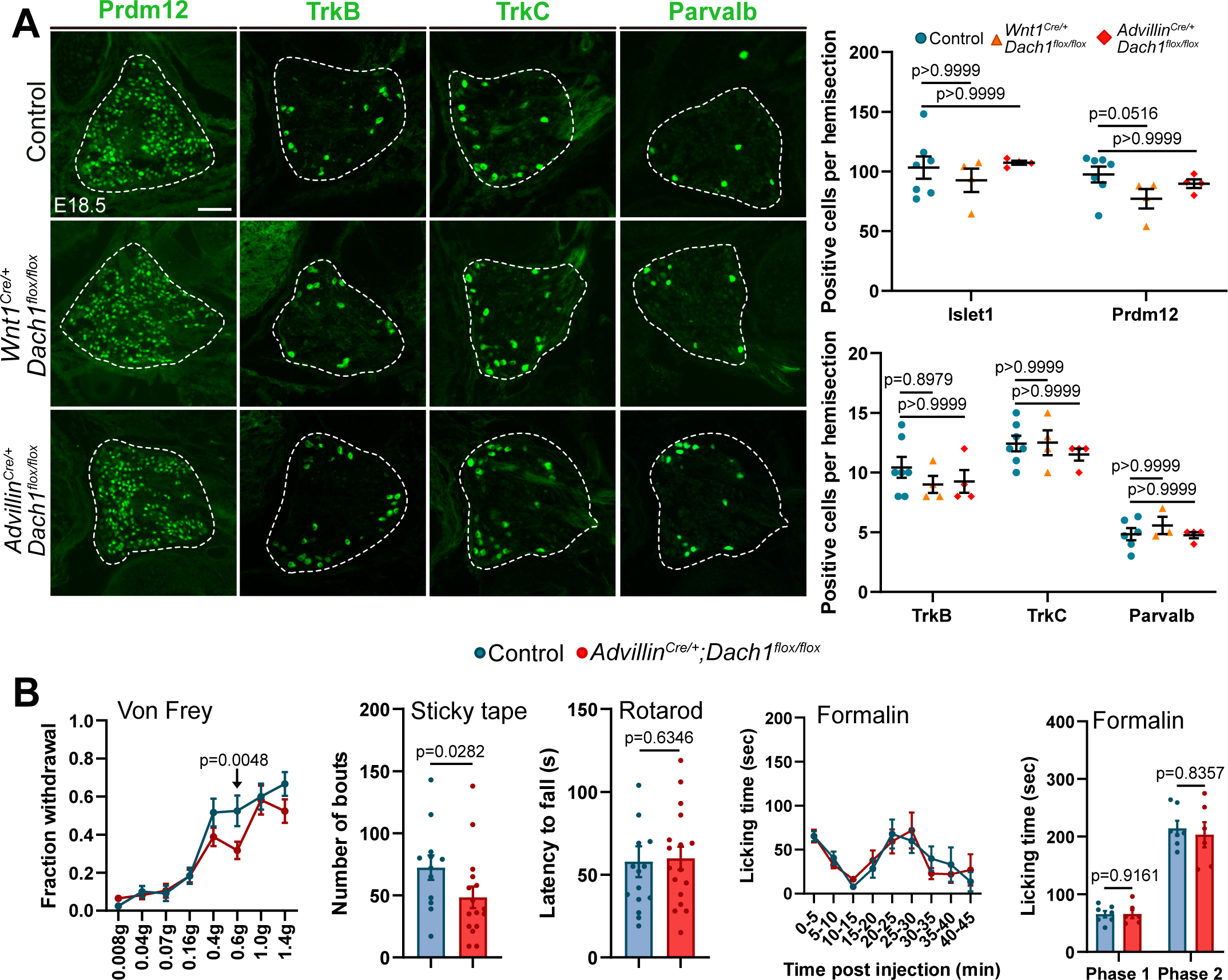
The loss of Dach1 in somatosensory neurons selectively impairs the sense of touch without affecting neurogenesis. **(A)** Left, Representative pictures of immunostaining targeting indicated somatosensory subtype markers performed on thoracic transverse DRG sections of control, *Wnt1^Cre/+^; Dach1^flox/flox^* (*WD1* cKO) *or Advillin^Cre/+^; Dach1^flox/flox^* (*AD1* cKO) E18.5 embryos. Right, Quantification analysis represented as scatter dot plot comparing the mean number of neurons immunostained for indicated marker on transverse hemisections of indicated genotypes. Each dot in this scatter plot indicates the mean value obtained for a single embryo. Graphical data in this panel and in the subsequent ones are presented as mean ± SEM. Kruskal-Wallis test with Dunn’s post hoc multiple comparison. **(B)** Behaviour analyses comparing control and *Advillin^Cre/+^; Dach1^flox/flox^* genotypes. Upper left, the response rate (fraction withdrawal) of control (n= 12) and *Advillin^Cre/+^; Dach1^flox/flox^* (n = 17) littermates to indicated forces applied to the plantar hindpaw using calibrated Von Frey filaments is indicated. Two-way ANOVA with Fisher’s LSD post hoc test. Upper middle, Quantification of the number of responses to the tactile stimulation of a sticky tape applied on the back skin of littermates of indicated genotypes. Mann-Whitney test. Upper right, Quantification of the time spent by littermates of indicated genotypes on an accelerating rotarod before falling. Mann-Whitney test. Lower left, Time course nocifensive response (licking time) of control (n = 7) and *Advillin^Cre/+^; Dach1^flox/flox^* (n = 6) littermates until 45 minutes following formalin injection. Two-way ANOVA with Fisher’s LSD post hoc test. Lower right, scatter dot plot indicating the response of individuals of indicated genotypes during the first (0 to 5 minutes following injection) and the second phase (15 to 45 minutes following injection). Mann-Whitney test. Dots in scatter plots indicate the measured value for a single individual. DRG in pictures are delineated by white dashed lines. Scale bars, 100 µm.

In adult *AD1* cKO mice, we also analysed histologically skin mechanosensory end organs innervation by A-LTMR^30^. This comprised analysis of LTMR endings innervating hair follicles, Meissner and Pacinian corpuscles and Merkel cells. No terminal morphology or innervation defects were observed in *AD1* cKO compared to control samples (**Figure S3A-E**). Thus, loss of Dach1 from its restricted period of expression does not result in peripheral innervation defects.

We next wanted to determine whether the loss of Dach1 affect the response of mice to tactile stimuli. We therefore performed Von Frey and sticky tape tests on *AD1cKO* and control mice. Despite their unaltered A-LTMR peripheral innervation, *AD1* cKO mice displayed reduced tactile sensitivity. We also performed Rotarod and Formalin tests, respectively to challenge kinesthesis and pain. No difference was observed between *AD1* cKO and controls indicating that loss of *Dach1* does not alter proprioception and nociception (**Figure 3B)**. Altogether, these results demonstrate that despite somatosensory neurons are appropriately generated in the absence of *Dach1*, its expression in A-LTMR is required for appropriate tactile sensitivity.

### The developmental loss of Dach1 primarily affects TrkB^+^ A-LTMR

To determine whether the transient broad expression of *Dach1* in developing somatosensory neurons reflects a transcriptional function aside from somatosensory neurogenesis, we performed a bulk RNA-sequencing of DRG harvested from E18.5 *WD1* cKO and littermate controls (**Supplemental Table 1**). Misregulated genes showed only modest fold changes, indicating that early depletion of *Dach1* has only discrete consequences on somatosensory neurons. Using a cut-off of log_2_(Fold Change) > |±0.15| with a p-value below 0.005, 30 genes were found to be differentially expressed between *WD1* cKO and controls, of which 16 and 14 were respectively reduced and increased in the *WD1* cKO embryos (**Figure 4A**). Despite these gene changes were evidenced at E18.5, analysis of the putative endogenous expression of downregulated genes at adulthood (using the online tool https://ernforsgroup.shinyapps.io/MouseDRGNeurons/)^9,17,18^ highlighted their prospective preferential enrichment in TrkB^+^ A-LTMR neuron classes (Aβ-RA-LTMR and Aδ-LTMR). This enrichment, correlated with the subsequent compartmentalized expression of Dach1, suggests its depletion during its broad expression period affects discrete subtypes of somatosensory neurons (**Figure 4B**).

**Figure 4.**
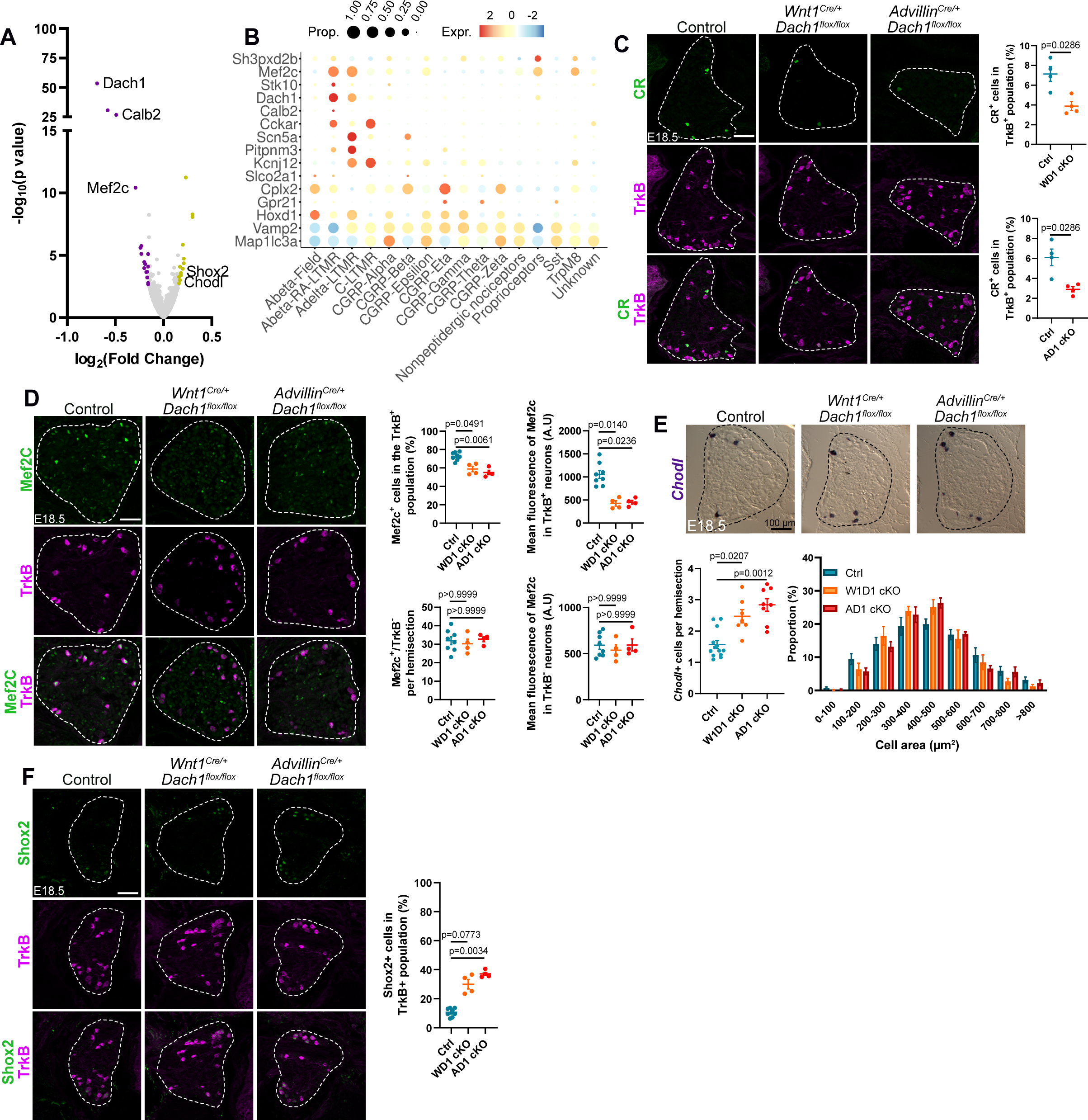
The loss of Dach1 selectively affects LTMR neurons. **(A)** Volcano plot showing the -log_10_(p-value) as a function of log_2_(Fold Change) of deregulated genes between DRG of E18.5 control (n = 5) and *WD1* cKO (n = 5) embryos. Genes were identified by bulk RNA-seq and selected based on a p-value < 0.005 and a log_2_(Fold Change) > |±0.15|. genes downregulated and upregulated above this threshold are respectively labelled in purple and yellow. **(B)** Bubble plot reporting the putative fraction of adult somatosensory subtypes that endogenously express the genes found misregulated in *WD1* cKO E18.5 DRG (generated using https://ernforsgroup.shinyapps.io/MouseDRGNeurons/)^9,17,18^. **(C)** Left, representative pictures of double immunohistochemistry targeting Calretinin (CR) and TrkB, performed on thoracic transverse DRG sections of control, *WD1* cKO and *AD1* cKO E18.5 embryos. Right, Quantification analysis of the mean percentage of CR^+^ neurons in the TrkB^+^ population in DRG of E18.5 embryos of indicated genotypes. Mann-Whitney tests. **(D)** Left, representative pictures of double immunohistochemistry targeting Mef2C and TrkB, performed on thoracic transverse DRG sections of control, *WD1* cKO and *AD1* cKO E18.5 embryos. Right, Quantification analyses of the mean percentage of Mef2C^+^ neurons inside or outside the TrkB^+^ compartment and quantification analyses of the mean fluorescence level (arbitrary unit, A.U.) of Mef2C signal inside or outside the TrkB^+^ compartment, in DRG of E18.5 embryos of indicated genotypes. Kruskal-Wallis tests with Dunn’s post hoc multiple comparison. **(E)** Upper panel, representative pictures of thoracic transverse DRG sections of control, *WD1* cKO and *AD1* cKO E18.5 embryos stained following *in situ* hybridization using a probe targeting *Chondrolectin* (*Chodl*) transcripts. Lower left, Quantification analysis comparing the mean number of neurons stained for *Chodl* on transverse hemisections of E18.5 embryos of indicated genotypes. Kruskal-Wallis test with Dunn’s post hoc multiple comparison. Lower right, histograms representing the percentage of *Chodl*-expressing cells of indicated range of soma area (µm²) observed in DRG sections of E18.5 embryos of indicated genotypes. **(F)** Left, representative pictures of double immunohistochemistry targeting Shox2 and TrkB, performed on thoracic transverse DRG sections of control, *WD1* cKO and *AD1* cKO E18.5 embryos. Right, Quantification analysis of the mean percentage of Shox2^+^ neurons in the TrkB^+^ population in DRG of E18.5 embryos of indicated genotypes. Kruskal-Wallis tests with Dunn’s post hoc multiple comparison. DRG in pictures are delineated by dashed lines. Scale bars, 100 µm.

Among the most downregulated genes found in our RNA-seq analysis were *Calb2*, a gene encoding for the neuronal modulator Calretinin (CR) found in Aβ-LTMR neurons innervating Pacinian corpuscles^31^ and *Mef2C*, a gene encoding for a transcription factor deeply involved in the developing central nervous system whose function remains unknown in the developing PNS^32,33^. The downregulation of proteins encoded by those genes was validated through co-immunostainings with TrkB at E18.5, comparing controls to both *WD1* cKO and *AD1* cKO conditions. In controls, CR was selectively found in a small population of TrkB-expressing neurons. Quantification of the percentage of CR^+^/TrkB^+^ neurons within the TrkB^+^ population confirmed its significant reduction in both cKO lines (**Figure 4C**). Mef2C similarly showed strong expression within the TrkB^+^ population but was also expressed in TrkB-negative neurons. Separate analysis of Mef2C expression inside and outside the TrkB^+^ compartment found no difference in the number of Mef2C^+^/TrkB^-^ cells and a modest but significant decrease of Mef2C^+^/TrkB^+^ neurons. Beside this reduction, the fluorescence signal of Mef2C in the remaining Mef2C^+^/TrkB^+^ neurons appeared consistently reduced in both cKO samples compared to controls. Quantification of the fluorescence level of this remaining Mef2C^+^ staining as a readout of Mef2C protein abundance in the nucleus confirmed its selective decrease in TrkB^+^ neurons (**Figure 4D**).

We then analysed the expression pattern of *Chodl* and *Shox2*, which were upregulated following *Dach1* depletion. *In situ* hybridization of *Chodl* revealed its endogenous restricted expression in a subset of large diameter somatosensory neurons, consistent with its previously reported expression in proprioceptors and in a subset of LTMR^34,35^. Analysis of *Chodl* revealed an increase in the number of *Chodl*-expressing cells in DRG of *WD1* cKO and *AD1* cKO E18.5 compared to controls (**Figure 4E**). To get clues of the type of neuron that start to ectopically express *Chodl* following *Dach1* depletion, we measured the diameter of the *Chodl*-expressing cells. We observed an enrichment in *Chodl*-expressing cells having a diameter between 300 and 500 µm^2^ in cKO compared to controls, suggesting that ectopic expression of *Chodl* selectively occurs in A-type large diameter neurons^6^. Finally, analysis of Shox2, a transcription factor endogenously found in a subset TrkB^+^ neurons, revealed an increased proportion of Shox2^+^/TrkB^+^ neurons in DRG of E18.5 *WD1* cKO and *AD1* cKO embryos compared to controls (**Figure 4F**).

Overall, these findings indicate that the loss of *Dach1* during its broad or restricted period primarily results in transcriptional defects associated with TrkB^+^ A-LTMR neurons. This together with its temporal expression dynamics establishes Dach1 as a *bona fide* broad-to-restricted transcription factor^9^. Since the transcriptional alterations observed when *Dach1* is depleted from the neural crest recapitulate those observed following its depletion in post-mitotic somatosensory neurons, they also suggest that the initiation of Dach1 expression is uncoupled from its functional turn-on.

### The temporal refinement of *Dach1* is a prerequisite for the appropriate transcriptional maturation of somatosensory subtypes

Depletion of *Dach1* during its broad expression period results in transcriptional defects clustered to the subpopulations in which it remains ultimately maintained. This observation prompted us to investigate the biological relevance of this temporal refinement. In other words, if Dach1 is anyway selectively involved in neurons in which it is maintained, is its repression essential in neurons in which it becomes turned off? To answer that question, we defined a strategy to maintain the pan-sensory expression of Dach1 by taking advantage of a previously described *Rosa26^lox-stop-lox-Dach1-IRES-EGFP^*mouse line which allows spatially and temporally controlled overexpression (OE) of *Dach1* following Cre-mediated excision^36^. We crossed this mouse line with *Advillin^Cre^* mice to generate *Advillin^Cre/+^; Rosa26^lox-stop-lox-Dach1-IRES-EGFP/+^* (*AD1* OE) embryos (**Figure S4A**). Pups of the desired genotype die around birth but otherwise develop until E18.5. At E13.5, a salt and pepper Dach1 expression pattern comparable to the control is observed (**Figure S4B-C**). Nevertheless, analysis of E18.5 *AD1* OE DRG revealed that Dach1 becomes effectively expressed in all somatosensory neurons later in development (**Figure S4D**). Thus, in *AD1* OE embryos, Dach1 is first appropriately refined before being broadly re-expressed at later stage (**Figure S4E**).

To start to characterize the consequences of the precluded temporal refinement of Dach1 in *AD1* OE embryos, we first quantified the number of Islet1^+^ neurons in DRG of *AD1* OE and control E18.5 embryos. We also analysed the proportion of neurons expressing Prdm12, TrkB and TrkC (**Figure 5A-B**). These analyses indicated that the different main subtypes of neurons are produced in appropriate numbers despite the pan-sensory maintenance of Dach1. Nevertheless, the possibility remains that Dach1 temporal refinement stays as a critical event for the subsequent steps of somatosensory neurons development. To further investigate this hypothesis, we collected DRG from E18.5 *AD1* OE and control embryos to perform a bulk RNA-seq analysis (**Supplemental Table 2**). Considering the selected differentially expressed genes following a cut-off of log_2_(Fold Change) > |±0.25| with an adjusted p-value below 0.005, 614 genes were found to be misregulated following *Dach1* forced pan-neuronal expression. 404 and 210 of them respectively showed reduced or increased expression, respectively. As expected, genes encoding for Prdm12, TrkA, TrkB or TrkC were not deregulated (**Figure 5C**).

**Figure 5.**
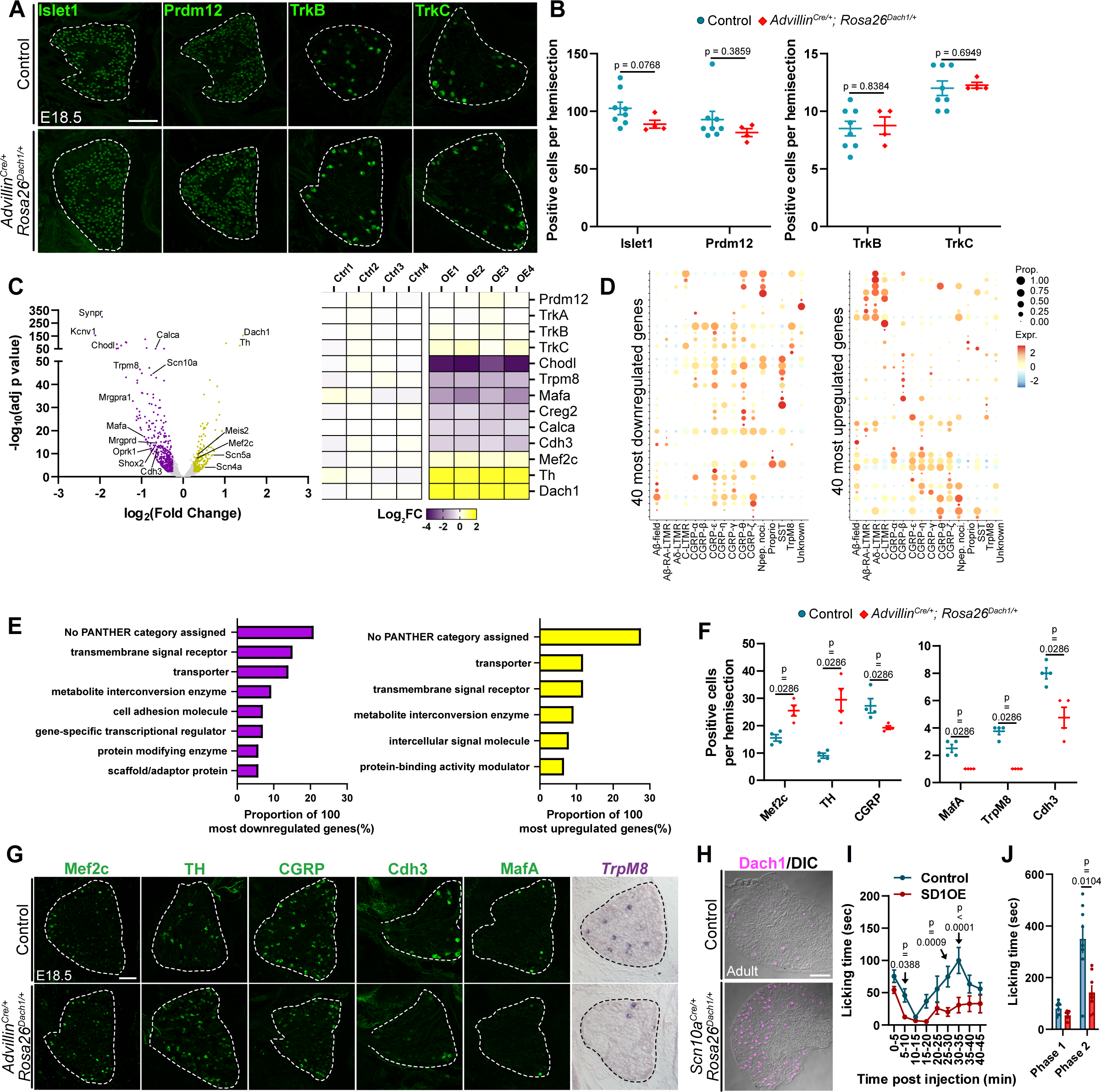
Countered temporal refinement of Dach1 in developing somatosensory neurons results in wide transcriptional maturation defects and altered nociception. **(A)** Representative pictures of immunostaining targeting indicated somatosensory subtype markers performed on thoracic transverse DRG sections of embryos of indicated genotypes. **(B)** Quantification analysis comparing the mean number of neurons immunostained for indicated markers on transverse hemisections of E18.5 of indicated genotypes. E18.5, Mann-Whitney tests. **(C)** Left, volcano plot showing the -log_10_(adjusted p-value) as a function of log_2_(Fold Change) of deregulated genes between DRG of E18.5 control (n = 4) and *Advillin^Cre/+^; Rosa26^lox-stop-lox-Dach1-IRES-EGFP/+^* (*AD1* OE; n = 4) embryos. Genes were identified by bulk RNA-seq and selected based on an adjusted p-value < 0.005 and a log_2_(Fold Change) > |±0.25|. Genes over this threshold which show downregulation in *AD1* OE are labelled in purple while the ones showing overexpression are labelled in yellow. Genes falling below the threshold are indicated in grey. Right, Heatmap of select genes showing their equivalent or differential expression between control and *AD1* OE conditions. **(D)** Bubble plot reporting the putative fraction of adult somatosensory subtypes that endogenously express the 40 most upregulated (left) or downregulated (right) genes identified in *AD1* OE E18.5 DRG (generated using https://ernforsgroup.shinyapps.io/MouseDRGNeurons/)^9,17,18^. **(E)** Gene ontology analysis reporting the protein class of the 100 most upregulated (left) or downregulated (right) genes identified in *AD1* OE E18.5 DRG. **(F)** Quantification analysis comparing the mean number of neurons stained for indicated markers on transverse hemisections of E18.5 embryos of indicated genotypes. Mann-Whitney tests. **(G)** Representative pictures of immunostainings with indicated antibodies and of *in situ* hybridization stainings using a probe targeting *TrpM8* transcripts performed on E18.5 thoracic DRG transverse sections of embryos of indicated genotypes. **(H)** Representative pictures of adult DRG transverse sections from control *Scn10a^Cre/+^; Rosa26^lox-stop-lox-Dach1-IRES-EGFP/+^* (*SD1* OE) mice immunostained for Dach1 and acquired with a differential interference contrast (DIC). **(I)** Time course nocifensive response (licking time) of control (n = 8) and *SD1* OE (n = 8) littermates until 45 minutes following formalin injection. Two-way ANOVA with Fisher’s LSD post hoc test. **(J)** Scatter dot plot showing the response of individuals of indicated genotypes during the first (0 to 5 minutes following injection) and the second (15 to 45 minutes following injection) phase of nocifensive behaviour of the formalin assay. Mann-Whitney tests. DRG in pictures are delineated by dashed lines. Scale bars, 100 µm.

Analysis of the putative endogenous expression of the 40 most upregulated and 40 most downregulated genes in adult somatosensory neurons revealed that these genes are differentially enriched. Many of the misregulated genes were characteristic of nociceptors, suggesting that this population is predominantly impaired following Dach1 forced pan-neuronal expression, while genes found in the endogenous Dach1 compartment were only found among upregulated genes, suggesting that ectopic expression of Dach1 results in ectopic expression of genes characteristic of its endogenous expression domain (**Figure 5D**). Gene ontology analysis of the 100 most upregulated and 100 most downregulated genes further revealed that they were enriched in terms correlating with transmembrane signal receptors or cellular transporters, reflecting their involvement in the functional components of somatosensory neurons (**Figure 5E**). Likewise, several of these misregulated genes were previously reported to have a physiological function in the activity of somatosensory neurons (*TrpM8*, *Scn10a, Mrgpra1, Oprk1*)^37–42^, or to be markers of defined somatosensory subpopulations (*Th*, *MrgprD*, *MafA*, *Calca*)^9,43^. Upregulation of *Th* and *Mef2C* and downregulation of C*alca*/CGRP, *MafA*, *TrpM8* and *Cdh3* were further validated by immunostaining or *in situ* hybridization on E18.5 transverse sections (**Figure 5F-G**). These results indicate that the sustained pan-sensory expression of Dach1 is not compatible with the appropriate transcriptional maturation of somatosensory neurons.

### Maintenance of Dach1 in nociceptors alters pain related behaviour

The defective transcriptional maturation resulting from the forced pan-sensory expression of Dach1 led us to hypothesize that the observed transcriptional changes could have repercussions on the physiological capacities of somatosensory neurons. Unfortunately, the postnatal death of *AD1* OE mice precluded further analysis of such somesthetic features. To overcome this obstacle, we seeked to cross *Rosa26^lox-stop-lox-Dach1-IRES-EGFP/+^* with a Cre-reporter mouse line allowing a narrower overexpression of Dach1 compared to the *Advillin^Cre^* mouse line. We used therefore the *Scn10a^Cre/+^* mice to generate *Scn10a^Cre/+^; Rosa26^lox-stop-lox-Dach1-IRES-EGFP/+^* (*SD1* OE) mice. *Scn10a* encodes for the nociceptor specific sodium channel Nav1.8 which is expressed from E14.5 in small diameter somatosensory neurons and thus allows the sustained expression of Dach1 in nociceptors^44^. *SD1 OE* mice were viable and fertile and showed wide expression of Dach1 in DRG (**Figure 5H**). Their nociception was evaluated using the formalin test. While the nocifensive response during the early phase was not markedly altered compared to controls, the response associated with the late phase was significantly reduced (**Figure 4G-H**). We thus conclude that the sustained expression of Dach1 in nociceptors is not compatible with a healthy sense of nociception.

## Discussion

### Dach1 as a new transcription factor required for appropriate behavioural reactivity to touch

Here, we report the previously unrecognized function of Dach1 in the developing somatosensory system. Dach1 is a transcription factor whose function has been extensively studied in other contexts and biological systems, notably in cancers and coronary angiogenesis^26,36,45,46^. Our results indicate that Dach1 is detected from somatosensory progenitors to mature post-mitotic somatosensory neurons. Initially pan-neuronal in developing DRG neurons, Dach1 becomes progressively turned off in most of these neurons to remain enriched in TrkB^+^ LTMR. In these neurons, Dach1 expression is required for appropriate behavioural reactivity to tactile stimulations. No alterations of mechanosensory fibers or of classical markers of LTMR subtypes are associated with embryonic Dach1 knock-out, suggesting that Dach1 is not involved in axonal growth or subtype specification. However, among the small set of genes downregulated following its depletion were genes directly involved in electrophysiological function (*Calb2*, *Scn5a*, *Kcnj12*, *Tmem266*). Also, genes overexpressed included several candidates encoding for cell-adhesion molecules (*Chodl*, *Pcdh7*, *Pcdh11x*) that could also affect the function of LTMR by modulating their peripheral and central ultrastructural features^47,48^.

### The broad-to-restricted expression of Dach1 is tightly linked to its functional turn-on

Recent single cell RNA-sequencing analyses of developing somatosensory neurons highlighted that neural crest cells biased to the somatosensory lineage first transition from an early immature post-mitotic neuronal state where several subtype-restricted transcription factors are initially expressed together. Among those are Runx1 and Runx3 which ultimately become mutually exclusive and are respectively involved in the specification of nociceptors and proprioceptors^9^. These broad-to-restricted transcription factors have in common that their depletion from their onset of expression results in defects restricted to the subtypes in which they are ultimately maintained. Here, we show that Dach1 behaves as a transcription factor with broad-to-restricted expression whose maintenance and extinction are both required to develop an unaltered sense of somesthesia. As such, we report that depletion of *Dach1* during its broad or restricted expression period predominantly alters gene expression in developing TrkB^+^ mechanosensory neurons. This observation argues for an uncoupling between the initiation of Dach1 expression in somatosensory progenitors and its functional turn-on in post-mitotic TrkB^+^ neurons. Indeed, genes found misregulated following depletion of *Dach1* during its broad expression period are equally found misregulated following its depletion when enriched in TrkB^+^ neurons. Moreover, despite the observation that Dach1 is already expressed in Sox10^+^ somatosensory progenitors, no significant function of Dach1 was found at this stage. Overall, these clues point out that the functional turn-on of Dach1 swiftly occurs following its expression refinement period in post-mitotic somatosensory neurons. Such temporal uncoupling between the expression of a transcription factor and its functional activity was previously highlighted during the lineage specification in the trunk neural crest. Environment may play a predominant role in this discrepancy by conditioning the availability of critical cofactors or regulatory proteins^49^. As such, Dach1 is found in dozens of cell types across the developing murine embryo^50^, suggesting that it may be engaged in multiple unrelated cell-type specification transcriptional programs. Indeed, while early reports concluded that *Dach1* KO mice die at birth with no obvious abnormalities^29,52^, subsequent analyses identifying a critical role for Dach1 in contexts as diverse as coronary artery development^26,36,46^, cortical somatostatin^+^ long-range projection neurons development^54^ or mammary gland development^56^ have highlighted the functional complexity of its contribution to diverse developmental processes. In such various already elucidated and putative functional involvements, the temporal repression of Dach1 where it could be deleterious rather than its induction whenever needed may represent an optimized way allowing it to be rapidly engaged in highly unrelated developmental processes defined by their own chromatin regulatory landscapes and active gene regulatory networks.

### The broad-to-restricted expression of Dach1 is physiologically relevant to produce functional somatosensory neurons

The progressive compartmentalization of broad-to-restricted transcription factors within the subtypes in which they ultimately play a function poses the question of the biological relevance of this progressive refinement. In other words, why do these transcription factors need to become subtype-specific if their reported early onset invalidation anyway results in subtype restricted defects? Opposite to broad-to-restricted transcription factors are transcription factors showing directly restricted late onset of expression. Among those is the ETS transcription factor Er81 whose neurotrophin-induced late post-mitotic expression is required for Ia proprioceptive neurons to synapse onto motor neurons in the spinal cord^51,53^. Experimentally challenging Er81 late onset of expression by inducing the precocious initiation of ETS signalling in early post-mitotic somatosensory neurons previously demonstrated that Er81 temporal sequence of expression matters during neuronal development. A developmental switch appears to operate during somatosensory neurons development which results in precocious post-mitotic ETS signalling being detrimental while late onset ETS signalling being mandatory for neuronal development^8^. Here, we challenged a similar paradigm by countering the temporal refinement of Dach1 through a Dach1 overexpression mouse line allowing its conditional expression upon Cre-mediated activity^36^. Through this strategy, we were able to sustain a pan-sensory expression of Dach1 during its late refined embryonic period or to express *Dach1* back into late post-mitotic *Scn10a*-expressing nociceptors. Forcing the maintenance of Dach1 broad expression resulted in wide transcriptional defects notably marked by the downregulation of a series of genes known to play functions in thermo/nociceptor (*TrpM8*, *Calca, Kcnv1, Oprk1, Scn10a*) and impaired the nocifensive behavioural response to formalin administration. These results demonstrate the importance to compartmentalize the expression of Dach1 in developing post-mitotic neurons and further reflect that its functional activation may result from a temporal developmental switch during the process of somatosensory neurogenesis. Altogether, these observations argue that the expression of Dach1 must become subtype restricted as soon as this transcription factor finds itself in an environment amenable to trigger its functional activity (**Figure S5**). This activity-dependent temporal refinement paradigm may serve as a basis to explain the expression dynamics of broad-to-restricted transcription factors.

### Molecular mechanisms involved in the broad-to-restricted expression of Dach1

What are the molecular cues resolving the selection of subtype-restricted transcription factors initially broadly expressed? It was previously shown that the NGF signalling pathway plays a critical role in refining the expression of several broad-to-restricted transcription factors, demonstrating that extrinsic cues are important for this refinement^1,9^. As Dach1 becomes progressively excluded from nociceptors, which represent an average 80% of DRG somatosensory neurons, we challenged whether the NGF signalling pathway is also required to resolve the subtype-restricted refinement of Dach1. However, we found that this mechanism does not apply to *Dach1*, which becomes also restricted in NGF^-/-^;Bax^-/-^ DRG. The observation that Prdm12, a major determinant of nociceptor lineage, remains expressed in NGF^-/-^;Bax^-/-^ DRG prompted us to hypothesize that it could support the temporal extinction of Dach1 in developing nociceptors. Prdm12 is a transcriptional regulator putatively acting as an epigenetic repressor. Increasing evidence indicate that it plays temporal stepwise functions from the emergence of the nociceptive lineage to mature nociceptors in adult, functionally switching its transcriptional targets as development proceeds. In early born nociceptors, Prdm12 represses the out-of-context expression of visceral determinants Phox2a/b and is required for nociceptor development and survival through the initiation and maintenance of the expression of *Ntrk1*/TrkA. In adult nociceptors, it modulates genes in a way that affects nociceptors homeostasis^20,22,23,25,55^. Through depletion of Prdm12 in late post-mitotic nociceptors using *Scn10a^Cre/+^; Prdm12^flox/flox^* mice, we were able to deplete Prdm12 in a model that did not result in Phox2a/b ectopic expression or in agenesis of the nociceptive lineage^25^. In the absence of Prdm12, Dach1 expression was expanded, indicating that Prdm12 contributes to its temporal refinement. Moreover, our ChIP experiments allowed us to identify a Prdm12-bound region in the Dach1 locus, suggesting that Dach1 is a direct transcriptional target of Prdm12. Intriguingly, the early pan-neuronal expression of Dach1 implies that in early born nociceptors, Dach1 and Prdm12 expression initially co-exist. Once again, this observation points out to a developmental switch where Prdm12 becomes competent to repress Dach1 at a time when its function is expected to become incompatible with the maturation of nociceptors.

In conclusion, through its maintenance in LTMR and extinction in nociceptors, Dach1 is critical for somatosensory neurons transcriptional maturation. The temporal refinement of its expression further coincides with the triggering of its functional competence, hypothesized to be driven by a change in the cellular context. Overall, our study demonstrates that the broad-to-restricted temporal expression followed by several transcription factors is physiologically relevant to achieve appropriate transcriptional maturation in somatosensory neurons.

## Acknowledgements

We thank members of the Bellefroid lab and Dr. Alexandre Pattyn for discussions and feedback about this manuscript. We a are grateful to Dr. Maya Dannawi and Dr. Panagiotis Tsimpos for help with the PCR screening of the Prdm12 knock-in mouse line. We thank the staffs of the CMMI (Center for Microscopy and Molecular Imaging) microscopy facility and of the IBMM-ULB Animal facility for their support. We are grateful to Dr. E. Bourinet and Dr. A. Pattyn for providing mouse lines and to Dr. Carmen Birchmeier for sharing MafA and c-Maf antibodies. This work was supported by grants from the FNRS (PDR T.0020.20 and T.0012.22) and by a grant from the International Brachet Foundation (IBS, project 22-3) to E.J.B. T.S. is supported by a grant from the Fondation Rose et Jean Hoguet and a ULB-UMons co-financed fellowship. S.D. is a FRS-FNRS postdoctoral fellow.

## Author Contributions

T.S., E.J.B. and S.D. conceived the study. L.R., E.J.B. and S.D. supervised the study. A.S collected and analysed ChIP-seq and ChIP-qPCR data with help from A.A. Y.A. generated the *Prdm12^KI^* mouse line. T.S., A.S. and S.K. contributed to the characterization of the *Prdm12^KI^* mouse line. T.S. and S.D. performed the histological and behaviour experiments. M.S. and G.S.-R. performed the RNA-sequencing and raw analysis of the data. K.R.-H. provided Dach1 transgenic mouse lines. S.D. prepared the figures and wrote the manuscript with inputs from all authors.

## Material and Methods

### Ressource availability

#### Lead contact

Further information and requests for resources and reagents should be directed to and will be fulfilled by the Lead Contact, Simon Desiderio.

#### Materials availability

This study did not generate new unique reagents or mouse lines.

### Experimental model and subject details

#### Mice

All mice were maintained on a C57Bl/6J background exc*ept Dach1^flox/flox^*and *Rosa26^Dach1^* mice which were of mixed C57Bl/6J and CD1 backgrounds. Mice were provided *ad libitum* with standard mouse lab pellet food and water and housed at room temperature with a 12h light/dark cycle. The experimental protocols were approved by the CEBEA (Comité d’éthique et du bien être animal) of the IBMM-ULB and conformed to the European guidelines on the ethical care and use of animals. *Dach1^flox/flox^* ^26^ and *Rosa26^Dach1^* ^36^ mouse lines were provided by Pr. Kristy Red-Horse (Department of Biology, Stanford University, Stanford, USA). *Advillin^Cre^* ^57^ and *Wnt1-Cre* ^58^ *mouse lines* were provided by Dr. P. Carrol and Dr. A. Pattyn (Institut des Neurosciences de Montpellier, Montpellier, France). The *Scn10a^Cre^*^44^ mouse line was provided by Dr. Emmanuel Bourinet (Institut de Génomique Fonctionnelle, Montpellier, France). The *Prdm12^flox/flox^*line was generated in our laboratory from a previous study^20^. *Bax^+/-^* (JAX:002994)^59^ and *NGF^+/-^* (JAX:003312)^60^ mice were ordered through Jackson Laboratories. The *Dach1^flox^* allele was detected by the primers forward 5’-CTCCTGAAGATGAGGAGCTCACCC-3’ and reverse 5′-AACAATTCTTGTCCTTCACGTGCCC-3’. The *Rosa26^Dach1^* allele was detected by the primers 5’-CTCTGCTGCCTCCTGGCTTCT-3’, 5’-CGAGGCGGATCACAAGCAATA-3’ and 5’-TCAATGGGCGGGGGTCGTT-3’. The Cre allele was detected by the primers forward 5’-CGATGCAACGAGTGATGAGGTTC-3’ and reverse 5’-GCACGTTCACCGGCATCAAC-3’. The *NGF KO* allele was detected by the primers 5’-CAGGCAGAACCGTACACAGA-3’, 5’-CCTTCTATCGCCTTCTTGACG-3’ and 5’-CTGTCACTCGGGCAGCTATT-3’. The *Bax KO* allele was detected by the primers 5′-GCAGAGGGTTAAAAGCAAGG-3′, 5′-CTTCCTGACTAGGGGAGGAG-3 and 5′-ACCCAGCCACCCTGGTCT-3′. The *Prdm12^flox^* allele was detected by the primers forward 5′-GCTGATCGAGTCCAGGAGAC-3′ and reverse 5′-CCAAACATCCACAACCTTCA-3′. The *Prdm12^KI^* allele was detected by primers forward 5’-GGTATCCACCCTTGTCTCCA-3’ and reverse 5’-CTTCAGCCCCTTGTTGAATACG-3’.

### Methods details

#### Tissue processing and fixation for Immunofluorescence

Whole embryos (E10.5-E15.5), vertebral column (E16.5, E18.5), DRG (adult), hindpaws and back skin were dissected in ice cold PBS and fixed for 10 to 15 minutes in 4% Paraformaldehyde at 4°C. Tissues were then rinsed with ice cold PBS and cryopreserved by overnight incubation in a solution of PBS containing 30% sucrose at 4°C and subsequently embedded in 15% sucrose-7,5% gelatin. For embryos, transverse sections of 12µm or 14µm, 25 µm for hindpaws and 30 µm for back skin, were done using a cryostat (Leica) and sections were kept at -20°C until use.

Slides were blocked in a solution of PBS with 0,1% Triton X-100 containing 10% of donkey serum for at least two hours at room temperature, then incubated in primary antibodies diluted in blocking serum overnight at 4°C. Slides were incubated in secondary antibodies diluted in PBST for two hours at room temperature. To reveal TrkB^+^ lanceolate endings in adult hairy skin, appropriate primary antibodies were instead incubated overnight at room temperature with no other modification to the protocol.

Primary antibodies used were: rabbit anti-Dach1 (1:500, Proteintech, 10914-1-AP), mouse anti-Islet1 (1:200, DSHB, 39.4D5), guinea pig anti-PRDM12 (1:1000)^20^, goat anti-TrkB (1:500, R&D System, AF1494), goat anti-TrkC (1:500, R&D System, AF1404), goat anti-TrkA (1:500, R&D System, AF761), guinea pig anti-Parvalbumin (1:500, Synaptic Systems, 195 004), guinea pig anti-c-Maf (1:10000, gift from Carmen Birchmeier), guinea pig anti-MafA (1:10000, gift from Carmen Birchmeier), rabbit anti-Th (1:250, Abcam, ab6211), sheep anti-Th (1:2000, Novus Biologicals, NB300-110), goat anti-Ret (1:50, R&D systems, AF482), goat anti-calcitonin gene related peptide (CGRP) (1: 200, Abcam, ab36001), rabbit anti-cleaved Caspase-3 (1:1000, Cell Signaling, #9661), rabbit anti-Mef2c (1:400, Cell Signaling, D80C1), rabbit anti-S100β (1:1000, Agilent, GA50461-2), chicken anti-NF200 (1:1000, Abcam, ab4680), rabbit anti-Calretinin (Millipore, AB5054, 1/1000), rabbit anti-Calretinin (Swant, 7697, 1/2000), mouse anti-Shox2 (1:400, Santa Cruz, sc-81955). Secondary antibodies used were: goat anti-rabbit Alexa Fluor 488 (1:1000, Invitrogen A11008), donkey anti-rabbit Alexa Fluor 488 (1:800, Abcam, ab150073), donkey anti-mouse Alexa Fluor 594 (1:1000, Invitrogen, A21203), goat anti-guinea pig Alexa Fluor 594 (Invitrogen, A11076), donkey anti-goat Alexa Fluor 594 (1:1000, Invitrogen, A11058), donkey anti-guinea pig Alexa Fluor 488 (1:800, Bio connect) and donkey anti-goat Alexa 405 (1:1000, Invitrogen, A21207). Anti-Shox2 primary antibody was directly detected using FlexAble Coralite ® Plus 488 antibody Labeling Kit for Mouse IgG2a (Proteintech, KFA041).

Images collection and analysis were performed using a widefield fluorescence microscope Zeiss Axio Observer Z1, a laser-scanning confocal microscope Zeiss LSM 710 using the Zeiss Zen black microscopy software. Image analysis was performed using ImageJ. For expression profile experiments cell counts were conducted with at least 10 DRG sections analysed per animal with at least two biological replicates. For comparisons of controls with mutant embryos, 18 to 24 DRG sections were analysed with at least four biological replicates. Samples from at least one control and one mutant animal were systematically collected on a same glass slide to reduce variability along the subsequent experimental procedures.

#### Wholemount Pacinian corpuscules immunostaining and quantification

Immunostaining of Pacinian corpuscules associated with the ulna was performed on dissected skinned forearms (radius and ulna) subsequently grossly trimmed from their muscles and connective tissue using a scalpel blade. To avoid damaging Pacinian corpuscules in the process, radius and ulna were kept together during the immunostaining procedure. Once the immunostaining was achieved the radius and ulna were separated and excess remaining surrounding tissue was carefully removed using forceps under an Olympus SZX16 binocular equipped with a X-Cite 120Q fluorescence illuminator. The wholemount immunostaining procedure was performed as described elsewhere for skin samples^61^. The number of Pacinian corpuscules associated with the anterior half of the ulna were counted in four different individuals of each genotype. The area of Pacinian corpuscules was measured using ImageJ.

#### Embryo processing and tissue fixation for *In situ* hybridization

Samples were fixed overnight in 4% Paraformaldehyde, rinsed with PBS and cryoprotected through overnight incubation in 30% sucrose at 4°C. Tissues were embedded and cut as described above. To process with the *in situ* hybridization, tissues were post-fixed in 4% PFA at room temperature for 20 min. After a 3 minutes-long proteinase K treatment, the slides were post-fixed in PFA 4% - 0,2% glutaraldehyde for 20 min. Tissues were then incubated in prehybridization solution (50% Formamide, 5X SSC, 50 µg/ml yeast extract, 1% SDS, 50 µg/ml heparin) for 2-3 hours at 70 °C. Slides were incubated overnight at 70°C with digoxigenin labelled riboprobes diluted in prehybridization solution. The next day slides were washed three times with a solution containing 50% Formamide, 5X SSC, 1% SDS), then three times with a solution containing 50% Formamide and 2X SSC. After a wash using TBS - 0,1% Triton X-100 (TBST), slides were blocked with 5% lamb serum diluted in TBST and incubated with an anti-digoxigenin antibody conjugated to Alkaline phosphatase (1/2000, Roche, 11093274910) overnight at 4°C. Slides were incubated twice in Alkaline phosphatase buffer (100mM Tris pH9.5, 100mM NaCl, 50mM MgCl2, 0.1% Tween 20) and incubated with the NBT and BCIP substrates (Roche) diluted in the same buffer. After PBS washes and dehydration, the slides were mounted with Eukitt hardening mounting medium (Sigma).

The *Chodl probe was* cloned by PCR from *Mus musculus* adult cortex cDNA using the following primers: *Chodl* forward 5′-CTGGTTTGGAACATGCTGGGC-3′ and *Chodl* reverse 5′-TTACTCCAAGATGTCTAAGTATACTGGTGG-3′. The PCR product was then cloned into the TOPO II dual promoter vector (Promega) and the <u>riboprobe</u> synthesized (linearized with SpeI, transcribed with T7). The plasmid containing the *TrpM8* probe was described elsewhere^62^.

Images collection was performed using a Zeiss Axioskop 2 equipped with a Axiocam 305 color camera and the ZEN blue software. Image analysis was performed with Image J. For comparisons of controls with mutant embryos, 18 to 24 DRG sections were analysed with at least four biological replicates. Samples from at least one control and one mutant animal were systematically collected on a same glass slide to reduce variability along the subsequent experimental procedures.

#### RNAScope

For RNAScope, we used fresh frozen individual DRGs from adult wild type mice, embedded in OCT (VWR cat.# 361603E), sectioned at 10 µm. Sections were kept at -80 °C until use. RNAScope multiplex fluorescent *in situ* hybridization (version 2) was carried out following the manufacturer’s protocol (Advanced Cell Diagnostics, ACD). Slides were removed from -80°C and were immediately post-fixed in cold (4°C) 4% paraformaldehyde for 15 min. The tissues were then dehydrated in 50% EtOH (5min), 70% EtOH (5 min) and 100% EtOH (2 X 5min) at room temperature. After air drying the slides, hydrophobic boundaries were drawn around each section with a hydrophobic pen. After a 10 min incubation with hydrogen peroxidase at room temperature, the tissues were subjected to a protease IV treatment for 25 minutes. Dach1-C1 (ACD Biotechne, cat# 412071) and Colq-C2 (ACD Biotechne, cat# 496211-C2) probes diluted (50:1:1 dilution, as directed by ACD due to stock concentrations) were added to the slides and hybridized for 2 hours at 40 °C. Signal amplification was carried out by adding AMP1, AMP2 reagents, each for 30 min at 40°C and AMP3 reagent for 15 minutes at 40 °C. Signal was developed in channel 1 with Opal570 (1/1500) and channel 2 with Opal 520 (1/1500). Slides were counterstained with DAPI and mounted with Dako Fluorescent Mounting Medium. Images were taken on a Zeiss LSM 710 confocal microscope and analysed with Image J.

### Generation of the *Prdm12^KI^* mouse line

The *Prdm12* knock-in (*Prdm12^KI^*) mouse line was generated through a CRIPSR-Cas9 mediated homology directed repair (HDR) strategy. A crRNA was designed using CRISPRdirect (http://crispr.dbcls.jp/) with the intent to produce a double strand break in the fifth exon of Prdm12, in 5’ from the STOP codon. 2.4 pmol/µl of annealed crRNA (10nM, Integrated DNA Technologies, 5’-GGCGCTCACAGCACCATGGC-3’) and the tracrRNA were injected with 200 ng/µl of recombinant Cas9 protein (Integrated DNA Technologies) and 10 ng/µl of a single-stranded DNA template (Megamer® single-stranded DNA fragment, GenScript) into the pronucleus of B6D2F2 zygotes which were subsequently transferred to pseudopregnant CD1 mice. The single-stranded DNA template was constituted (from 5’ to 3’) of: a *Prdm12 exon 5* homology arm (75 nucleotides), a GGTGGT Glycine linker sequence, three repeated Flag tag sequences, one V5 tag sequence terminated by a TGA stop codon, one Internal Ribosome Entry Site (IRES), one sequence encoding for a Venus fluorescent protein, one TGA stop codon followed by a 3’UTR homology arm (75 nucleotides). Founder mice having properly integrated the *Prdm12^KI^* allele were screened by PCR and appropriate integration of the knockin-in region was validated by Sanger sequencing. Germline transmission of the selected *Prdm12^KI^* founder mouse was then validated by PCR.

#### Chromatin immunoprecipitation and data processing

The spinal cord with attached DRG of E12.5 embryos (10 embryos/condition) or isolated DRG of E13.5 (18 embryos/condition) were collected in ice cold PBS and fixed in 1% Formaldehyde (Thermo scientific) in Phosphate Buffer Saline (PBS, pH 7.5) for 15 minutes at room temperature in a shaking plate. The samples were quenched by adding a final concentration of 125 mM Glycine (Sigma Aldrich) during 5 minutes at room temperature. Finally, the samples were washed twice in ice cold PBS and stored at -80°C. Tissues were resuspended in 900 µL of SDS Lysis buffer (50mM Tris pH 8.1, 10mM EDTA, 1% SDS, cOmplete EDTA-free Protease Inhibitor Cocktail, Roche, #cat 05892791001) and sonicated for four cycles of 10 minutes (30 seconds ON 30 seconds OFF) at high setting using a Bioruptor Plus sonication device (Diagenode). The resultant fragmented chromatin was centrifuged at 10.000 g for 10 minutes and the supernatant was collected. The chromatin was diluted two times in Chromatin dilution buffer (16.7mM Tris pH8.1, 2mM EDTA, 0.1% SDS (wt/vol), 1% Triton X100(vol/vol), 167mM NaCl, cOmplete EDTA-free Protease Inhibitor Cocktail). 10% of the chromatin was de-crosslinked for input and eluted in a total volume of 50µl. 9µL of de-crosslinked inputs were run on an agarose gel to determine chromatin shearing efficiency. The chromatin was confirmed to be between 150 and 400 bp before proceeding to the chromatin immunoprecipitation. 800 µl of chromatin was mixed with 3200 µl of chromatin dilution buffer and 5 µg of rabbit polyclonal anti-V5 (Abcam, ab15828) or normal rabbit IgG (Merk, 12-370). The samples were incubated at 4°C overnight. The next day, 30 µL of magnetic beads (Cell signaling, #cat 9906) were added and incubated in rotation at 4°C during 2h. Beads were collected using a magnetic stand and washed three times with low salt buffer (20mM Tris pH8.1, 2mM EDTA, 1% SDS (wt/vol), 200mM NaCl) for 5 minutes at 4°C in rotation and one time with high salt buffer (20mM Tris pH8.1, 2mM EDTA, 1% SDS (wt/vol), 0.5M NaCl) for 5 min at 4°C. The bound chromatin was subsequently eluted in 200µL of elution buffer (1%SDS (wt/vol), 100mM NaHCO3) at 65°C during 30 minutes. A magnetic stand was used to separate the beads and the supernatant containing chromatin was collected in a separate Eppendorf tube. Input and immunoprecipitated chromatin samples were brought to a final volume of 200µL and were de-crosslinked and the DNA was extracted using the following protocol. To each sample, a final concentration of 200mM NaCl was added and incubated overnight at 65°C. The next day, the samples were treated with RNAse (1µL/sample) for 30 minutes at 37°C. Followed by the addition of 8µL of Tris HCl pH 6.5, 4µL 0.5M EDTA and 1µL Proteinase K (20mg/mL) for 1h at 45°C. Finally, the DNA was extracted using the High Pure PCR Product Purification kit (Sigma Aldrich, REF 11732676001). The final chromatin was diluted in 50µL of elution buffer.

For ChIP-qPCR, the input DNA was diluted 100 times and each immunoprecipitated chromatin sample was diluted 10 times. The Luna Universal qPCR Master Mix following manufacturer’s instructions for qPCR, adding 5uL of DNA per reaction. The qPCR was runned on a StepOnePlus PCR instrument (Applied BioSystem). Fold enrichment was calculated as the percentage of input values. Primers forward 5’-AATGAGCCTTTGCTTCGCTT-3’ and reverse 5’-TTTTTCTCAGCCCATCTGCG-3’ were used to detect the intronic *Dach1* region bound by Prdm12. Primers forward 5’-GCCTCGCTGACCGAAACTAA -3’ and reverse 5’-CACCCTTGCCAGCTGTTTTG -3’ were used to detect the Prdm12-bound region close to *Dbx1*.

For ChIP-seq, the collected chromatin was sequenced using an Illumina Novaseq (BrightCore Platform, http://www.brightcore.be/) Paired-end reads were trimmed using TrimmomaticPE to remove Illumina universal adapters. For data analysis, trimmed reads were mapped to mouse genome mm10 using Bowtie2^63^ with default parameters for paired-end sequencing. Duplicate reads were removed with MarkDuplicates tools (Picard suite). Peaks were called with the callpeak tool from MACS2 package^64^ with following parameters: -f BAMPE -g mm -q 0.05 – nomodel –call-summits -B –SPMR. For visualisation, deeptools package was used to generate bigwig files^65^.

#### RNA sequencing and data processing

Dorsal root ganglia were dissected in cold RNAse Free PBS and stored at −80°C in TRIZOL (ThermoFisher cat.# 15596026). RNA was extracted using the illustra RNAspin Mini RNA isolation kit (GE Healthcare cat.# 25-0500-70). RNA quality was assessed by measuring the RNA integrity number (RIN) using a Fragment Analyzer HS Total RNA Kit. Library preparation for RNA-Seq was performed using the Illumina Stranded mRNA Prep, Ligation and the Illumina RNA UD Indexes Set A, Ligation starting from 300 ng of total RNA. The size range of the final cDNA libraries was determined by applying the SS NGS Fragment 1- to 6000-bp Kit on the Fragment Analyzer (average 340 bp). Accurate quantification of cDNA libraries was performed by using the QuantiFluor™ dsDNA System (Promega). cDNA libraries were amplified and sequenced by using an S4 flow cell NovaSeq6000; 300 cycles, 25 Mio reads/sample from Illumina. Sequence images were transformed with BaseCaller Illumina software to BCL files and demultiplexed to fastq files with bcl2fastq v2.20.0.422. Sequencing quality was determined using FastQC v. 0.11.5 software (http://www.bioinformatics.babraham.ac.uk/projects/fastqc/).

Sequences were aligned to the *Mus musculus* GRCm39 genome using the STAR aligner software version 2.7.8a, allowing for 2 mismatches within 50 bases. Subsequently, read counting was performed using featureCounts version 2.0.1. Read counts were analyzed in the R/Bioconductor environment version 4.1.0 (www.bioconductor.org) using the DESeq2 package version 1.32.0. Candidate genes were selected as those with a FDR-corrected P-value < 0.05 and absolute log2 fold-change >1. Genes were annotated using the M musculus GTF file mm10 version 105 used to quantify the reads within genes. RNA-seq data have been deposited at Gene Expression Omnibus (GEO) under accession numbers GSE270844 and GSE270845. Gene ontology analysis was performed using PANTHER^66^ and selecting for terms associated with Protein Component.

#### Quantification of Meissner corpuscules density

Immunofluorescence with anti-S100β and NF200 antibodies was carried out on 25 µm sections of glabrous pedal pads from hindpaws to visualize Meissner corpuscules. The number of Meissner corpuscles in a section was counted and the area of the epidermis of the same section was traced and measuread using Image J. Density of Meissner was determined by diving the number of corpuscles by the area of the epidermis.

#### Quantification of hair follicles associated LTMR endings

Images of serial sections of back hairy skin were cryosectioned at 30 µm thickness. Quantification of the relative proportion of innervated hair follicles was performed using ImageJ by counting the number of hair shafts associated with NF200^+^ or TrkB^+^ endings on four independent hairy skin sections of 16 mm length. Counterstaining with S100β allowed visualisation of terminal Schwann cells associated with LTMR terminals to facilitate the quantification.

#### Quantification of Merkel cells innervation

Merkel cells innervation was visualized through Cytokeratine K8 (CK8) and NF200 immunostaining in both hairy and glabrous skin on sections of 30 or 25 µm thickness respectively. Confocal images were processed using ImageJ and CK8^+^ Merkel cells were considered innervated if touched by a NF200^+^ nerve fiber.

#### Quantification of relative fluorescence level

Quantifications of the relative fluorescence level of Dach1 and Mef2C immunostainings were performed in ImageJ. By tracing a line crossing the major axis of individual nuclei and measuring the resulting mean gray value of fluorescence intensity of nucleus pixels crossing this line, the average Dach1^+^ of Mef2C^+^ signal intensity was manually calculated for every cell analysed. Analysed images were obtained using a laser-scanning confocal microscope Zeiss LSM 710 and the Zeiss Zen black microscopy software. Care was taken to maintain the exact same parameters of image acquisition for all experiments. Whenever appropriate, Superforst glass slides on which samples were collected always contained side-by-side serial sections of one control and one experimental condition to minimize variation during the subsequent immunostaining procedures.

#### Behavioral assays

Unless otherwise specified, all tests were performed on mixed cohorts of males and females. All testing were performed by experimenters blind to genotypes. Details about the sex of mice and their exact genotypes can be found in **Supplemental Table 3**.

##### Von Frey

Mice were placed on an elevated wire mesh grid into PVC chambers. Before the test, mice were habituated to the device for 1 hr for two consecutive days. On the testing day, mice were placed in the chamber 1 hr before Von Frey filaments application. The test was performed as previously described^67^. During the test, withdrawal response following Von Frey filament application on the palm of the left hind paw was measured. Starting with the lowest force, each filament ranging from 0.008 g to 1.4 g was applied 10 times in a row with a break of 30 s following the fifth application. During each application, bend filament was maintained for 4–5 s. The number of paw withdrawals for each filament was counted.

##### Sticky Tape

A 2 cm^2^ of laboratory tape was placed on the upper-back skin of mice just before they were placed on an elevated wire mesh grid into PVC chambers. The number of tape-directed reactions was then counted during 5 min. Considered responses were body shaking like a ‘wet dog’, hindlimb scratching directed to the tape, trying to reach the tape with the snout and grooming of the neck with forepaws.

##### Rotarod

Rotarod behavioural experiment was carried out over a 5-day period. Mice were trained on the Rotarod apparatus at a fixed speed of 10 rotations per minute (rpm) on the first day and tested for the next four days. On the days of the tests, the speed of the device was progressively increased from 10 to 40 rpm over a five-minute period. The latency to fall of each mouse was then recorded and the mean time was reported after three consecutive trials separated by a recovery period of 5 minutes.

##### Formalin

Mice were habituated for 15 minutes on the day of the test in a PVC chamber with 3 mirrors on the walls and wire mesh grid on the floor. The dorsal part of the left hindpaw of the mouse was then injected with 10 µL of a 5% formalin solution and directly observed during a 45-minute period. The licking time of the injected paw was recorded during 5 minutes intervals. Formalin assay including AD1 cKO animals was selectively performed on male cohorts while the assay including SD1 OE mice was performed on both males and females.

### Quantification and statistical analysis

Quantitative analyses were carried out on at least 3 independent animals of each genotype. All statistical analyses and the generation of graphs were performed using the GraphPad Prism software. Statistical tests as well as the number of cells or animals per group used in each experiment are described in the legends of the figures and supplementary figures wherever appropriate. In case a dataset did not follow normal distribution, nonparametric statistical tests have been applied. In case a dataset did not contain enough biological replicates to be able to determine whether they follow a normal distribution or not, non-parametric tests have been also applied. No statistical methods have been used to determine sample sizes. All data are presented as the mean ± the standard error of the mean (SEM). Exact p-values are reported wherever appropriate.

