## Supplementary Figures 1 to 5 for "The temporal refinement of *Dach1* is a key step in the functional maturation of primary somatosensory neurons"

### Supplemental figure 1

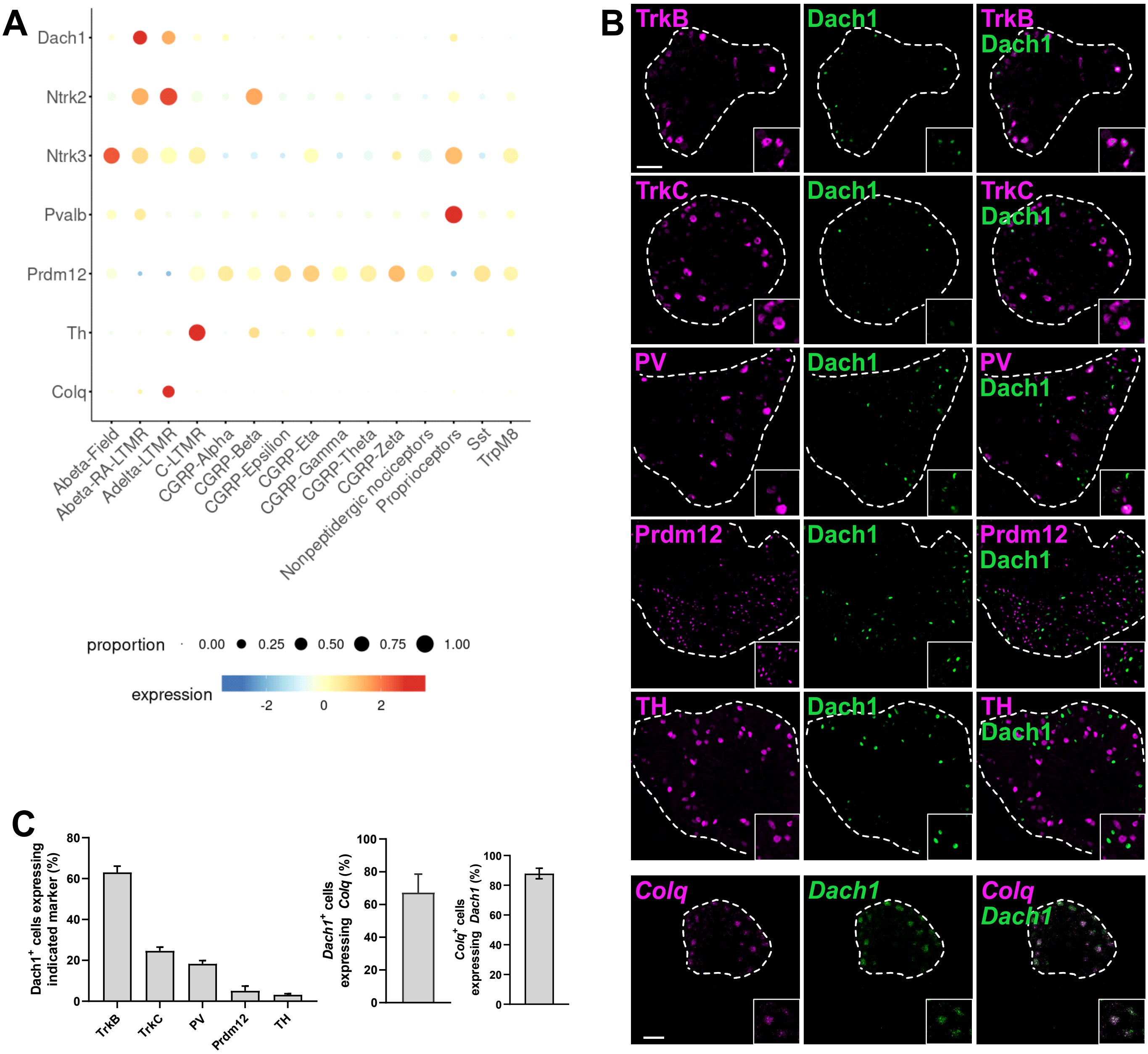


**Supplemental Figure 1. Dach1 is predominantly retained in TrkB⁺ somatosensory neurons at adulthood.**

**(A)** Bubble plot of *Dach1* and selected genes highlighting *Dach1* enriched expression in Aβ-RA-LTMR and Aδ-LTMR in adult somatosensory neurons (generated using https://ernforsgroup.shinyapps.io/MouseDRGNeurons/).

**(B)** Double immunohistochemistry targeting Dach1 and indicated somatosensory subtype markers performed on transverse sections of thoracic dorsal root ganglia harvested from adult wild-type mice. Co-staining of *Colq* and *Dach1* was performed by fluorescent *in situ* hybridization (RNAscope). Scale bar, 100 µm.

**(C)** Percentage of colocalization of Dach1 and indicated neuronal markers (N= 2, mean ± SEM)

### Supplemental figure 2

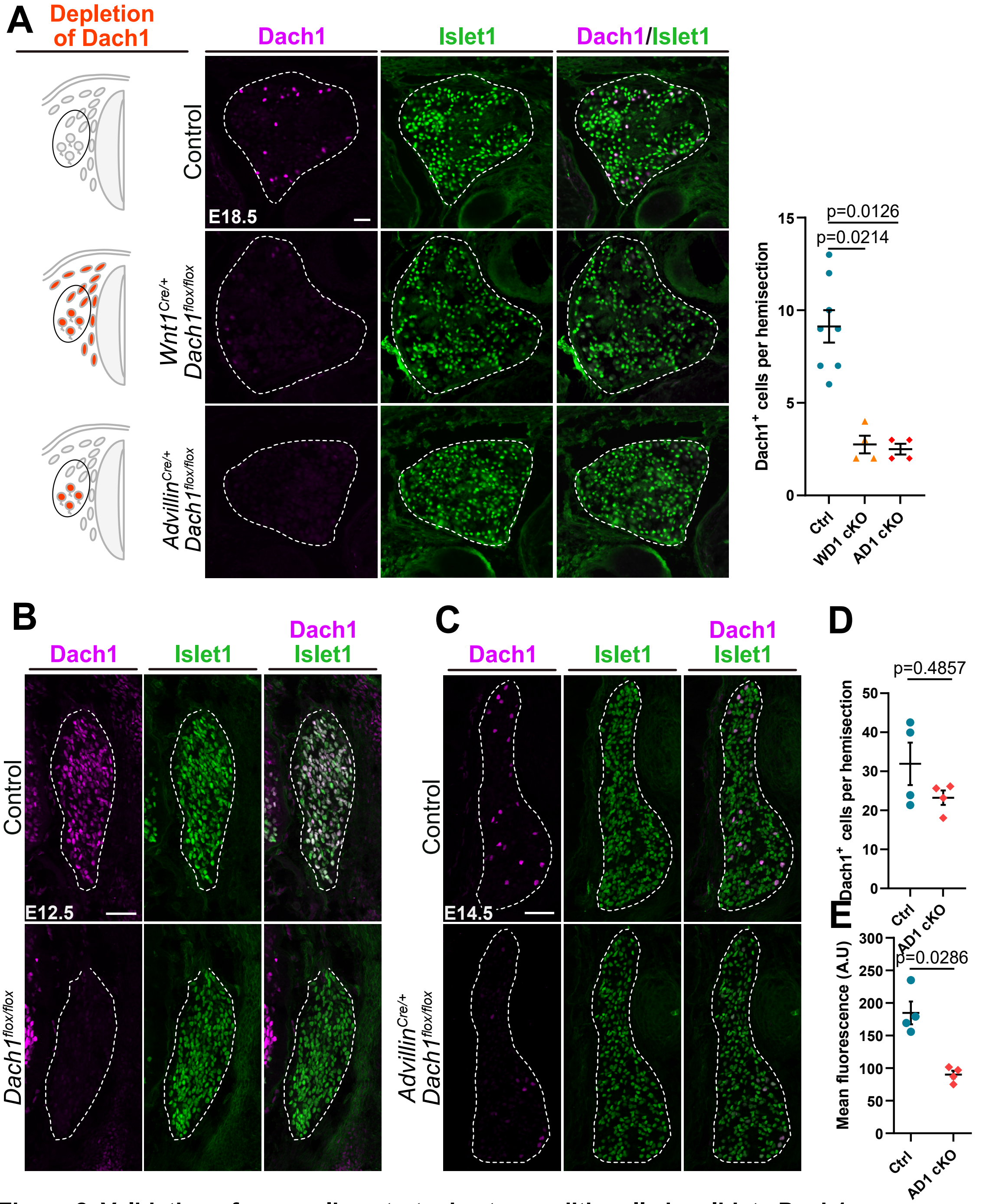


**Supplemental Figure 2. Validation of mouse line strategies to conditionally invalidate Dach1.**

**(A)** Left, schematic representation of the two strategies established to deplete Dach1 from the neural crest or from post-mitotic somatosensory neurons. Cells expected to be targeted by each startegy are labelled in orange. Middle, Representative pictures of double immunostainings targeting Dach1 and the pan-sensory neuron marker Islet1 performed on thoracic transverse DRG sections of control, *Wnt1*$^{Cre/+}$;*Dach1*$^{flox/flox}$ (*WD1* cKO) and *Advillin*$^{Cre/+}$;*Dach1*$^{flox/flox}$ (*AD1* cKO) E18.5 embryos. Right, quantification analysis represented as scatter dot plot comparing the mean number of neurons immunostained for Dach1 detected in dorsal root ganglia on transverse hemisections of indicated genotypes. Each dot in this scatter plot indicates the mean value obtained for a single embryo. Graphical data in this panel and in the subsequent ones are presented as mean ± SEM. Kruskal-Wallis test with Dunn's post hoc multiple comparison.

(**B**) Representative pictures of double immunostainings targeting Dach1 and the pan-sensory neuron marker Islet1 performed on thoracic transverse DRG sections of control and *Wnt1*$^{Cre/+}$;*Dach1*$^{flox/flox}$ (*WD1* cKO) E12.5 embryos. Note that Dach1 is already completely abolished at this stage in the *WD1* cKO line.

(**C**) Representative pictures of double immunostainings targeting Dach1 and the pan-sensory neuron marker Islet1 performed on thoracic transverse DRG sections of control and *Advillin1*$^{Cre/+}$;*Dach1*$^{flox/flox}$ (*AD1* cKO) E12.5 embryos. Note that Dach1 is still detected at this stage, depsite at a lower fluorescence level.

(**D**) Quantification of the mean number of Dach1$^+$ cells per DRG cells on hemisections of E14.5 control and *AD1* cKO embryos. Mann-Whitney test.

(**E**) Quantification of the mean fluorescence level of Dach1$^+$ immunostaining (arbitrary unit, A.U.) in DRG neurons of E14.5 control and *AD1* cKO embryos. Mann-Whitney test.

Dorsal root ganglia are delineated by white dashed lines.

Scale bar, 50 µm.

### Supplemental figure 3

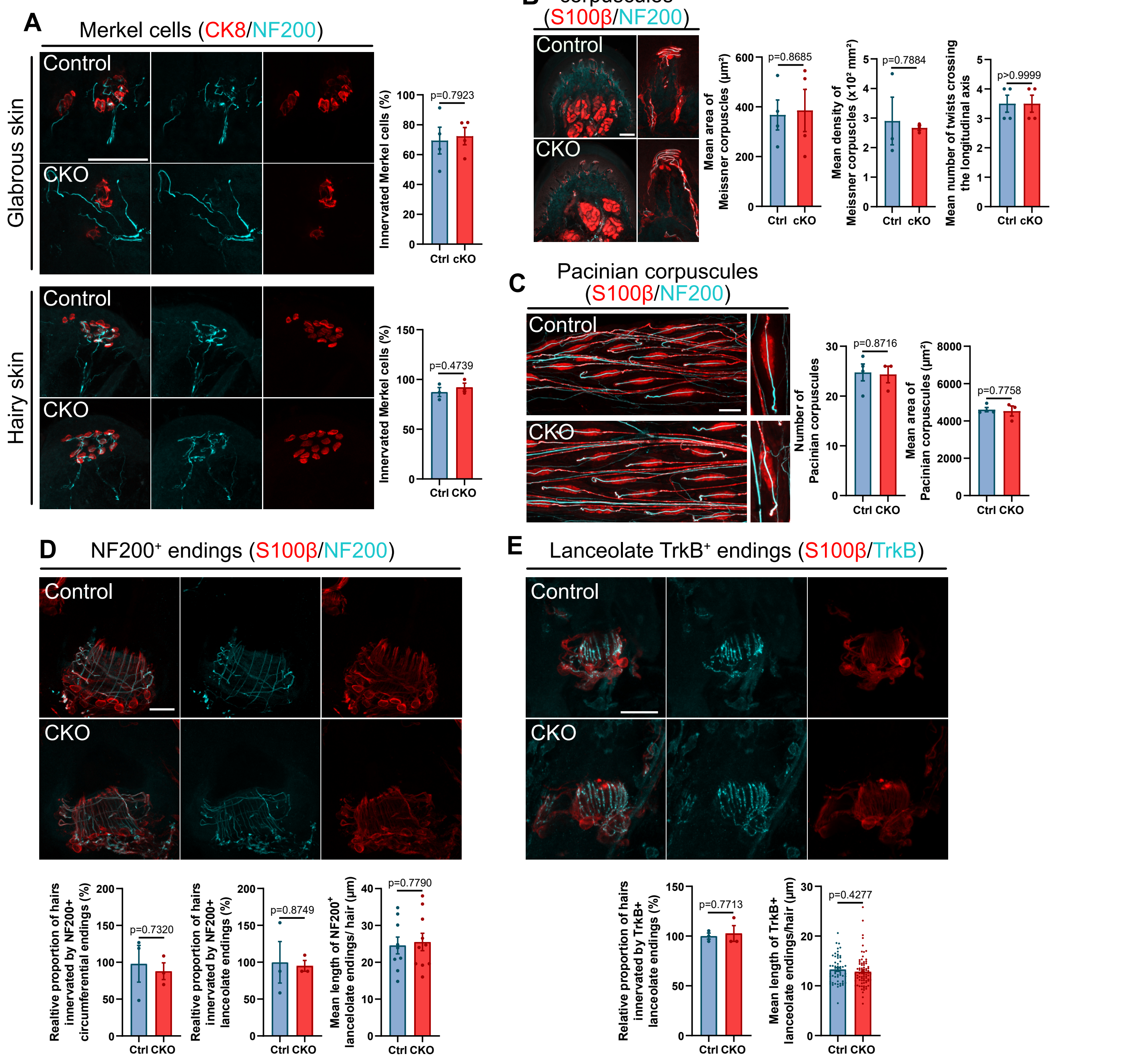


**Supplemental Figure 3. The loss of Dach1 does not affect the innervation and morphology of touch receptor end-organs.**

**(A)** Left, Representative immunostaining images of Merkel cells and their innervating fibers in glabrous or hairy skin cross-sections collected from mice of indicated genotypes. Right, Quantification of the proportion of innervated Merkel cells. Student's unpaired t tests. Each dot represents the mean percentage of innervated Merkel cells from a single individual. Scale bar, 50 µm.

**(B)**Left, Representative immunostaining images of Meissner corpuscules and their innervating fibers in glabrous skin cross-sections collected from mice of indicated genotypes. Right, Quantification of the area (left) and density (center) of Meissner corpuscules as well as the number of NF200+ fiber twists crossing the longitundinal axis (right) as a readout of their innervation complexity . Student's unpaired t tests. Each dot represents the mean value obtained for the indicated parameters in a single individual. Scale bar, 100 µm.

**(C)**Left, Representative whole-mount immunostaining images of Ulnar Pacinian corpuscules and their innervating fibers from mice of indicated genotypes. Right, Quantification of the number (left) and mean area (right) of Ulnar Pacinian corpuscules. Student's unpaired t tests. Each dot represents the mean value obtained for the indicated parameters in a single individual. Scale bar, 100 µm.

**(D)**Up, Representative immunostaining images of NF200+ Aβ circumferential and lanceolate endings innervating guard hair follicles and their S100β+ associated terminal Schwann cells. Bottom, Quantification of the relative proportion of guard hairs innervated by Aβ circumferential (left) or Aβ lanceolate endings (center) in *AD1* cKO mice compared to control. Each dot represents the mean value obtained for the indicated parameters in a single individual. Bottom right, Quantification of the mean length of Aβ lanceolate endings innervating a hair follicle in mice of indicated genotypes. Each dot represents the mean lenght of Aβ lanceolate endings innervating a single guard hair follicle. Student's unpaired t tests. Scale bar, 20 µm.

**(E)**Up, Representative immunostaining images of TrkB+ Aδ lanceolate endings innervating zigzag hair follicles and their S100β+ associated terminal Schwann cells. Bottom left, Quantification of the relative proportion of zigzag hairs innervated by Aδ longitudinal endings in *AD1* cKO mice compared to control. Each dot represents the mean value obtained for a single individual. Bottom right, Quantification of the mean length of Aδ lanceolate endings innervating a zigzag hair follicle in mice of indicated genotypes. Each dot represents the mean lenght valnnervating a single hair follicle. Student's unpaired t tests. Scale bar, 20 µm.

### Supplementary figure 4

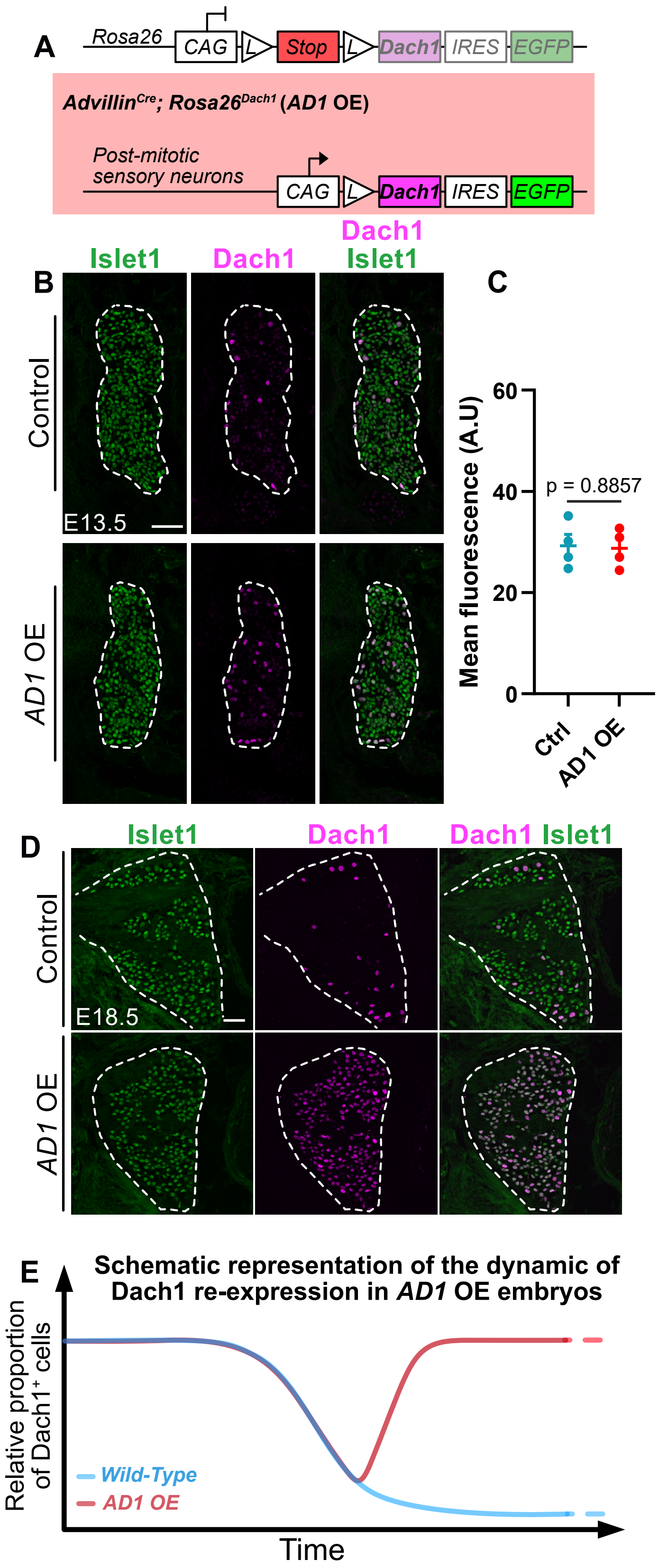


**Supplemental Figure 4. Transgenic mouse line strategy to perturb the temporal refinement of Dach1 *in vivo.***

**(A)** Schematic representation of the strategy established to force late stage pan-neuronal expression of Dach1 in post-mitotic somatosensory neurons (red background). To make reading easier, embryos overexpressing *Dach1* through the *Advillin$^{Cre}$ and Rosa26$^{Dach1}$* alleles are abbreviated *AD1* OE.

**(B)** Representative pictures of double immunostainings targeting Dach1 and the pan-sensory neuron marker Islet1 performed on thoracic transverse DRG sections of control and *AD1* OE E13.5 embryos. Scale bar, 50 µm.

**(C)** Quantification of the mean fluorescence level of Dach1 immunostaining (arbitrary unit, A.U.) among Islet1$^{+}$ DRG neurons of E13.5 control and *AD1* OE embryos. Mann-Whitney test.

**(D)** Representative pictures of double immunostainings targeting Dach1 and the pan-sensory neuron marker Islet1 performed on thoracic transverse DRG sections of control and *AD1* OE E18.5 embryos. Scale bar, 50 µm.

**(E)** Schematic representation of the characteristic features of the startegy established to counter the temporal refinement of Dach1. The blue line summarizes the dynamic of Dach1 broad-to-restricted expression observed in wild-type embryos. The red line recapitulates the results observed in the *AD1* OE embryos, namely that *Dach1* temporal restriction is initially properly triggered at E13.5 before being broadly expressed again at later stage.

**Supplemental figure 5**

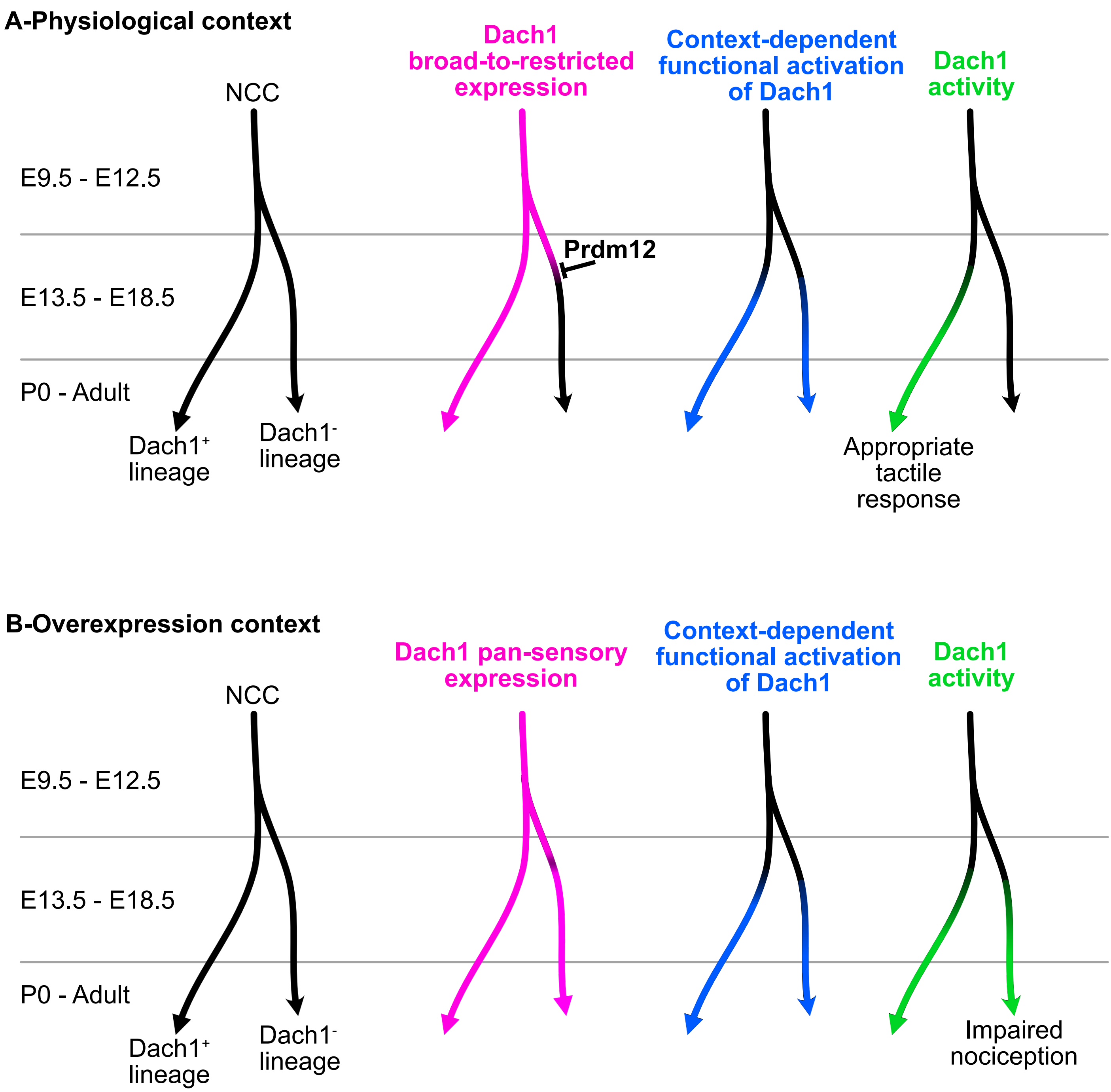


**Supplemental Figure 5. Model of Dach1 refinement and functional activity.** Dach1 is expressed following a broad-to-restricted expression dynamic, being first expressed in all developing somatosensory precursors before seggregating into a branch representing neuronal subtypes maintaining Dach1 expression and an other branch representing neuronal subtypes repressing Dach1, in part through the activity of Prdm12 (A-Pink). This change timely occurs before a contextual developmental switch driving the functional activation of Dach1 (A-Blue). The combination of its timely refined expression and triggering of its functional competence results in an appropriate subtype-restricted Dach1 activity required for appropriate tactile response (A-Green). The forced maintenance of Dach1 in the putative Dach1$^{-}$ lineage (B-Pink) associated with the triggering of its functional activation (B-Blue) consequently results in Dach1 becoming functional in the putative Dach1$^{-}$ lineage, ultimately resulting in impaired nociception (B-Green).
